# AGO1 in neural progenitor cells orchestrates brain development and sociability via LIN28A-REELIN axis

**DOI:** 10.1101/2025.10.01.679670

**Authors:** Hyunsu Do, Dayeon Kim, Roberto Jappelli, Hyeokjun Yang, Yiseon Jeon, Geurim Son, Mingyu Ju, Jihoon Son, Insook Ahn, Christina K Lim, Jungeun Ji, Jihae Lee, Young-Kook Kim, Joon-Yong An, Seung-Hee Lee, Daeyoup Lee, Fred H. Gage, Jinju Han

**Affiliations:** Graduate School of Medical Science and Engineering, Korea Advanced Institute of Science and Technology (KAIST), Daejeon, 34141, Republic of Korea; Laboratory of Genetics, Salk Institute for Biological Studies, La Jolla, CA 92037, USA; Department of Biological Sciences, KAIST, Daejeon, 34141, Republic of Korea; Department of Integrated Biomedical and Life Science, Korea University, Seoul, 02841, Republic of Korea; L-HOPE Program for Community-Based Total Learning Health Systems, Korea University, Seoul, 02841, Republic of Korea; School of Biosystem and Biomedical Science, College of Health Science, Korea University, Seoul, 02841, Republic of Korea; Department of Biochemistry, Chonnam National University Medical School, Jeollanam-do, 58128, Republic of Korea; Center for Synaptic Brain Dysfunctions, Institute for Basic Science, Daejeon, 34141, Republic of Korea; BioMedical Research Center, KAIST, Daejeon, 34141, Republic of Korea; KAIST Stem Cell Center, KAIST, Daejeon, 34141, Republic of Korea

## Abstract

AGO1, an essential RNA binding protein (RBP) in RNA interference, is associated with autism spectrum disorder (ASD). However, the precise functions of AGO1 in brain development and related disorders remain largely unexplored. Here, we report the critical roles of AGO1 in neural progenitor cells (NPCs) in promoting sociability and shaping the structure of the developing brain. *Ago1* knockout (KO) in the mouse brain leads to hyposociability, a characteristic symptom of ASD. In human forebrain organoids, *AGO1* KO disrupts the formation of ventricle-like structures and delays cortical layer development. We discovered that *AGO1* KO results in a loss of polarity in NPCs, which subsequently reduces neuronal development. These *AGO1* KO phenotypes originate from the failure to suppress LIN28A in NPCs. AGO1 is primarily localized in the nucleus within NPCs and binds to the LIN28A promoter region, thereby inhibiting LIN28A transcription. We found that increased LIN28A in *AGO1* KO NPCs reduces REELIN expression, a key regulator of brain development, by binding directly to the REELIN mRNA. The aberrant polarity phenotype in *AGO1* KO NPCs was successfully ameliorated by either LIN28A knockdown or recombinant REELIN treatment. Collectively, our findings elucidate the intricate molecular mechanisms orchestrated by nuclear AGO1 in NPCs and underscores its pivotal roles in brain development, providing significant insights into ASD.

## Introduction

RBPs interact with RNAs to form a ribonucleoprotein complex, playing a vital role in regulating gene expression at both transcriptional and post-transcriptional levels^1–7^. The majority of RBPs, characterized by their RNA binding domains, are ubiquitously expressed across different tissues, indicating their fundamental importance in a wide range of biological processes. The specific biological roles of RBPs are influenced by the RNA repertoire present in each tissue. RBPs are particularly critical in brain development and function, as the brain expresses a much more diverse RNA populations compared to other tissues^8^. Furthermore, the association of dysfunctional RBPs with various neurological diseases highlights the biological significance of RBPs in the brain^9^.

ASD is a prominent condition categorized among the neurodevelopmental disorders (NDD). While the etiology of ASD is multifaceted and complex, genetic variations have been identified as major causative factors in the development of ASD^10^. Large-scale genomic studies have identified high-confidence ASD risk genes, which are approximately 100 genes with significant associations to ASD^11–14^. These risk genes are involved in various pathways critical for cortical neurogenesis, with functions including chromatin regulation and synaptogenesis^12^. Notably, the risk genes significantly overlap with the target genes of RBFOX1^13^ or FMRP, which are both well-known RBPs^14^. In addition, a significant proportion of ASD risk genes—about 38% (449 genes) of the 1,162 annotated by SFARI (as of February 2024) - feature RNA binding motifs, based on RBP2GO database records^15–17^. This large number of RNA binding motifs underscores the potential role of RBPs in the development of ASD.

AGO1, an RBP belonging to the Argonaute (AGO) protein family, has been identified as a strong risk factor for ASD^18–24^. AGO family proteins involved in RNA interference primarily regulate gene expression post-transcriptionally in the cytoplasm^25,26^, but they also possess important nuclear functions like transcription regulation^27,28^. Numerous genetic variations in *AGO1* have been identified in the genomes of patients with ASD including developmental delay, cognitive deficits, and distinctive facial characteristics^23,29–36^. While other proteins in the AGO family are also implicated as ASD risk factors, AGO1 stands out due to the higher abundance of its genetic variations and the greater number of reported cases of patients with AGO1 variations compared to variations in other AGO proteins^34^. However, the specific molecular mechanisms by which *AGO1* variations contribute to NDD such as ASD remain unclear. To better understand the pathogenic effects of *AGO1* variations and their association with ASD-related behavioral symptoms and traits, an in-depth exploration of the functional mechanisms of AGO1 within the brain is essential.

In this study, we have elucidated the critical roles of AGO1 in brain development, particularly in regulating cortical structure, which is vital for sociability. We validated the link between AGO1 and ASD by showing that *AGO1* depletion in the mouse brain leads to hyposociability, a characteristic feature of ASD. Through *AGO1* KO in human embryonic stem cells (hESCs), we revealed that *AGO1* KO notably disrupts the structures of human forebrain organoids. The phenotypes of organoids were confirmed in two-dimensional (2D) cultured NPCs and neurons, revealing changes in NPC polarity and a delay in neuronal development. We identified the primary molecular function of AGO1 in the nuclei of NPCs, where AGO1 suppresses LIN28A transcription. LIN28A suppression subsequently enhanced the expression of genes translated through endoplasmic reticulum-associated pathways, such as REELIN. We found that LIN28A directly binds to REELIN mRNA and reduces its stability. The abnormal polarity in AGO1 KO NPCs was restored either by suppressing the upregulated LIN28A with shRNAs or by adding recombinant REELIN protein to compensate for the downregulated REELIN. Collectively, our findings unveil a novel molecular mechanism regulated by AGO1 in brain development and advance our understanding of the connection between AGO1 and ASD.

## Results

### *Ago1* KO in the brain leads to hyposociability

While variations in *AGO1* have been identified in the genomes of patients with ASD, it is not clearly known whether ASD symptoms are caused by dysfunctional AGO1 in the brain or by other factors. To address this question, we aimed to validate if the loss-of-function of *Ago1* in the brain results in ASD symptoms using a mouse model. For this, we generated brain-specific *Ago1* KO mice using *Nestin-Cre* transgenic mice (Fig. 1a). We were able to obtain an adequate number of *Ago1* conditional KO (cKO) mice unlike *Ago1* global KO mice, which are born with abnormal Mendelian ratios (about 9% of homozygous KO) (MGI: 5285081,5576271).

**Fig. 1:**
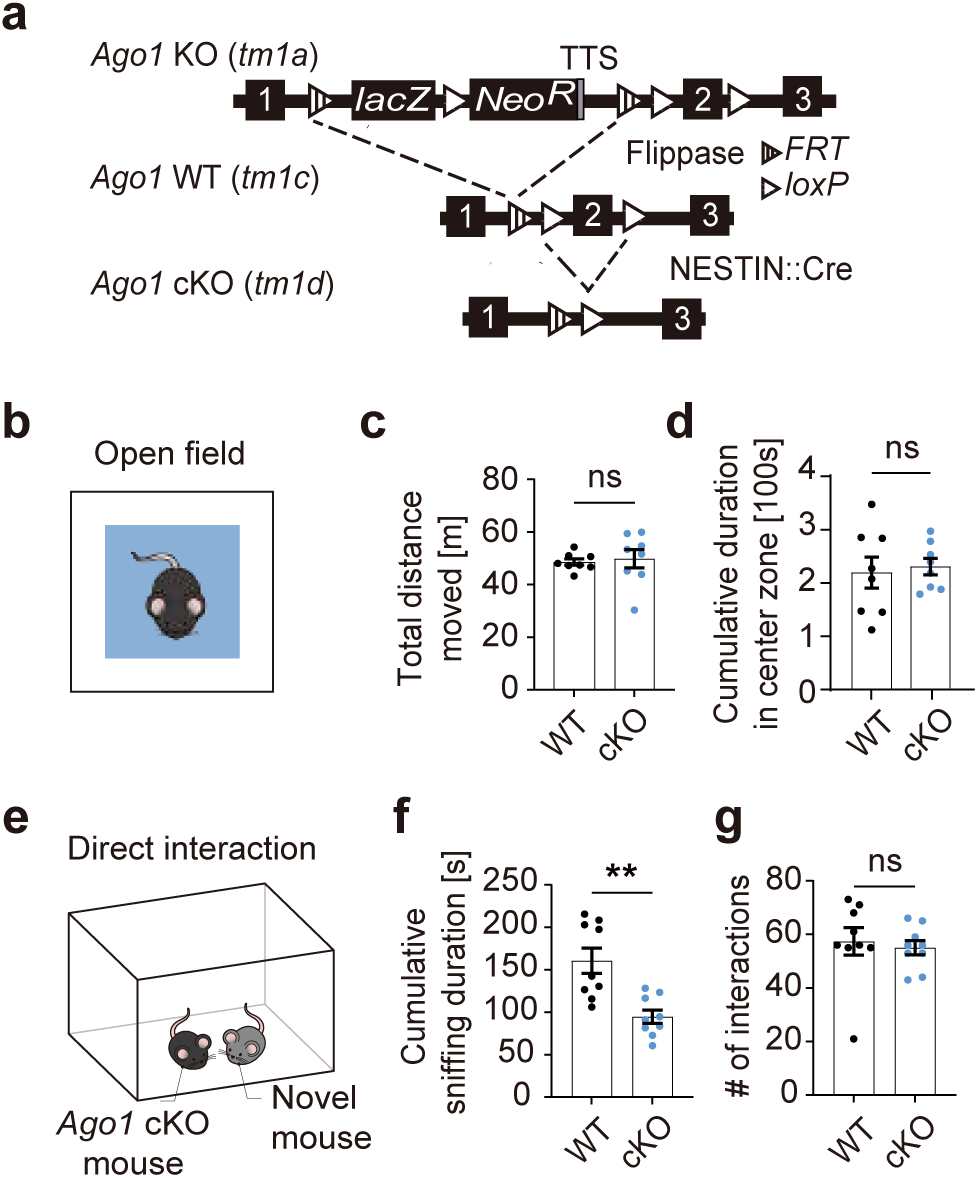
Behavioral assessments in *Ago1* cKO mice. a. Scheme of *Ago1* cKO strategy. TTS, transcription termination sequence. b. Diagram of the open field test. The center zone,defined as a 20cm x 20cm area in the center of the open chamber, is highlighted in pale blue (c, d) Tracking of locomotor activity in the open field test was conducted with 8 male mice each for both *Ago1* WT and cKO groups, with all activities recorded over a 15-minute period. c. Total moved distance (mean ± SEM, unpaired Student’s t-test*, p*=0.7414). d. Cumulative duration spent in center zone. (mean ± SEM, unpaired Student’s t-test*, p*=0.7446). e. Diagram of the direct interaction test with a novel mouse. (f, g) The direct interaction test was conducted with 9 male mice from both *Ago1* WT and cKO groups. Their interactions with a novel mouse were recorded over a 10-minute period. f. The duration of direct social sniffing (mean ± SEM, unpaired Student’s t-test, \*\**p*=0.0012). g. The frequency of direct interactions (mean ± SEM, unpaired Student’s t-test*, p*=0.6918).

To assess motor activities and anxiety levels in the *Ago1* cKO mice, we conducted an open field test, but there was no significant difference in total distance moved and cumulative duration spent in the center zone between *Ago1* cKO mice and control mice (Fig. 1b-d). To further clarify differences in sniffing behavior, we conducted a direct interaction test. The total number of interactions with a novel mouse was comparable between *Ago1* cKO and wild type (WT) mice. However, *Ago1* cKO mice exhibited significantly shorter cumulative sniffing duration towards the novel mouse (Fig. 1e-g). These data indicate that *Ago1* cKO mice are less engaged in social interaction, which is a representative feature of ASD, and suggests that dysfunctional AGO1 in the brain is sufficient to lead to the development of social deficits in mice, as has been shown in human patients.

### *AGO1* KO disrupts the structure of forebrain organoids and delays cortical development

To evaluate the effects of *AGO1* KO on human brain development, forebrain organoids were generated from hESCs after *AGO1* KO. For *AGO1* ablation in the hESC genome, the CRISPR/Cas9 genome editing technique was applied. We designed two distinct single guide RNAs (sgRNAs), each targeting a separate constitutive exon of *AGO1* (Fig. 2a). Following the introduction of sgRNAs and Cas9 into hESCs, we isolated a hESC colony derived from a single hESC. The colony containing indels in the *AGO1* gene was identified as *AGO1* KO hESCs. Additionally, for comparative control, we selected another hESC colony also originating from a single hESC that maintained an intact *AGO1* gene despite undergoing identical sgRNA/Cas9 transfection. The *AGO1* KO was confirmed through genomic DNA sequencing and Western blot analysis (Fig. 2b,c).

**Fig. 2:**
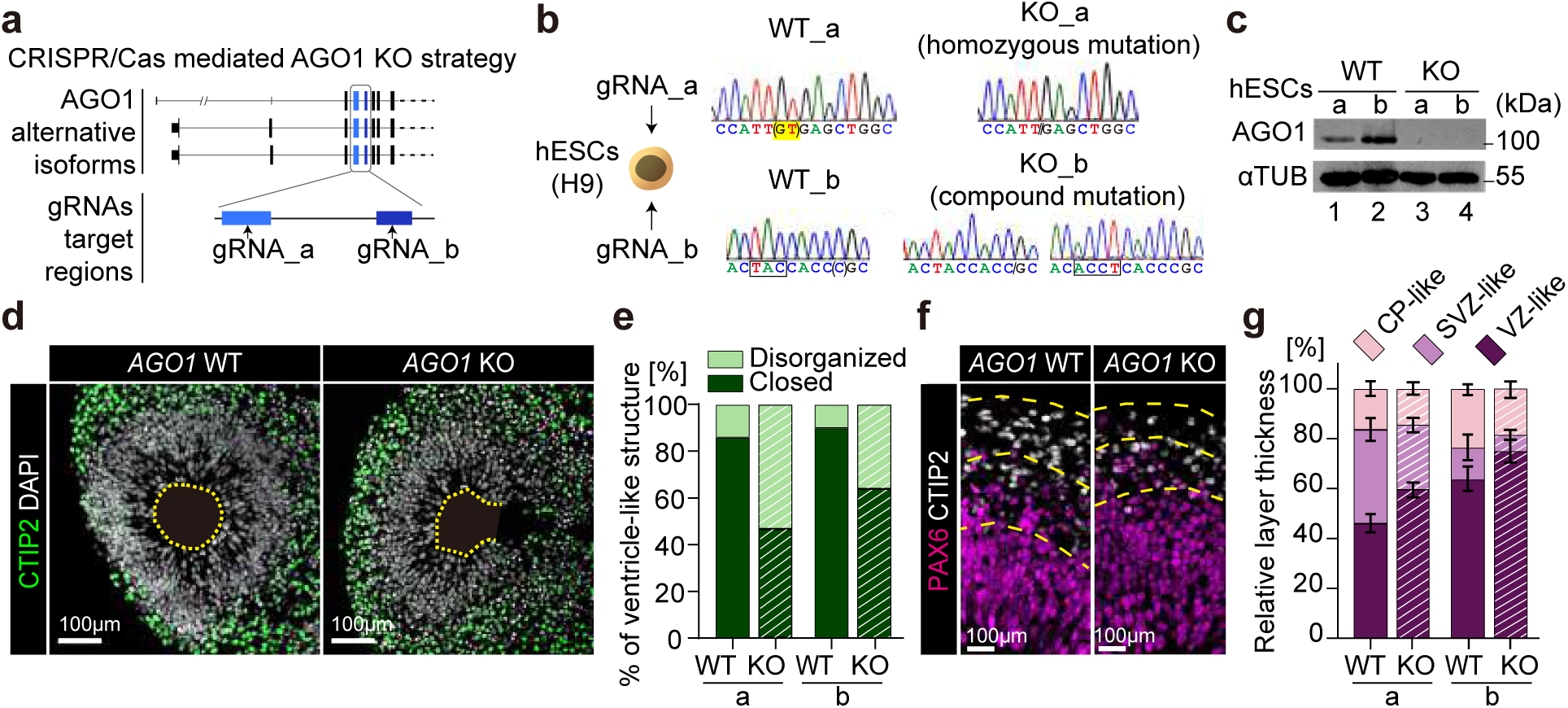
The effect of *AGO1* KO on human forebrain organoids. a. Strategy for generating *AGO1* KO hESCs using the CRISPR-Cas9 system. b. Sanger sequencing results displaying genome editing outcomes in *AGO1.* Presented are 2 different *AGO1* KO hESC clones carrying indels in *AGO1,* alongside 2 control hESC clones with intact *AGO1*. c. Validation of *AGO1* KO in hESCs via Western Blot. Human AGO1 protein (∼100 kDa) was analyzed with α-TUBULIN (αTUB; ∼55 kDa) as a control. d. Representative confocal images from a single plane of AGO1 KO and WT forebrain organoids, showing ventricle-like structures, quantified in (e). Day 50 forebrain organoids were stained with CTIP2 (green) for ventricle-like layers and DAPI (blue) for nuclei. Scale bar = 100 μm. e. Quantification of disorganized ventricle-like structures in day 50 forebrain organoids. (n=9 *AGO1* WT organoids, n=8 *AGO1* KO organoids). f. Representative confocal images from a single plane of forebrain organoids, used for analyzing layer thickness in (g). Day 50 organoids were stained for CTIP2 (white) as a ventricle-like layer marker and PAX6 (magenta) as a progenitor cell marker. The cortical plate (CP)-like layer is defined by CTIP2 expression, the subventricular zone (SVZ)-like layer by co-expression of CTIP2 and PAX6, and the ventricular zone (VZ)-like layer by PAX6 expression. Scale bar = 100 μm. g. Quantification of relative layer thickness in forebrain organoids at day 50, based on specific marker staining of ventricle-like structures (n=14 *AGO1* WT: n=11 WT_a, n=3 WT_b; n=17 *AGO1* KO: n=12 KO_a, n=5 KO_b, mean ± SEM).

The deletion of *AGO1* did not affect the overall size of the forebrain organoids during their development (Extended Data Fig. 1a,2b). However, notable malformations were observed in the ventricle-like structures within the *AGO1* KO organoids (Fig. 2d-g and Extended Data Fig.1c-f). In both *AGO1* KO organoids (KO_a and KO_b), approximately 50% of the ventricle-like structures displayed disorganized structures (Fig. 2d, e). Furthermore, we found that *AGO1* KO organoids showed alterations in thickness of cytoarchitecture, representing developing cortical layers. The ventricular zone (VZ)-like cell layer appeared thicker, whereas the subventricular zone (SVZ)-like cell layers were thinner in *AGO1* KO organoids compared to controls, suggesting a developmental delay in *AGO1* KO forebrain organoids (Fig. 2f,g and Extended Data Fig.1d-f).

### *AGO1* KO influences cell adhesion and neuronal development

To elucidate the biological processes resulting in the phenotypes observed in *AGO1* KO forebrain organoids, we used NPCs and neurons derived from hESCs through the 2D cell culture method (Extended Data Fig.2a). These cells are relatively homogeneous compared to organoids, thereby facilitating the investigation of the effects of *AGO1* KO and exploration of molecular mechanisms at specific stages of neuronal development. No significant effect of *AGO1* KO on NPC production was observed, as evidenced by the consistent number of cells immunostained with the NPC marker proteins SOX2 and NESTIN (Fig. 3a and Extended Data Fig.3b). Additionally, analysis of cell cycle dynamics revealed that *AGO1* KO did not influence symmetric cell division among NPCs (Fig. 3b). Moreover, comparable numbers of TUJ1- and MAP2-positive cells were observed in both *AGO1* KO and WT neurons upon NPC differentiation (Fig. 3c and Extended Data Fig.3c). These findings suggest that *AGO1* does not significantly affect the fate determination of ESCs into NPCs and neurons; neither does it impact NPC proliferation.

**Fig. 3:**
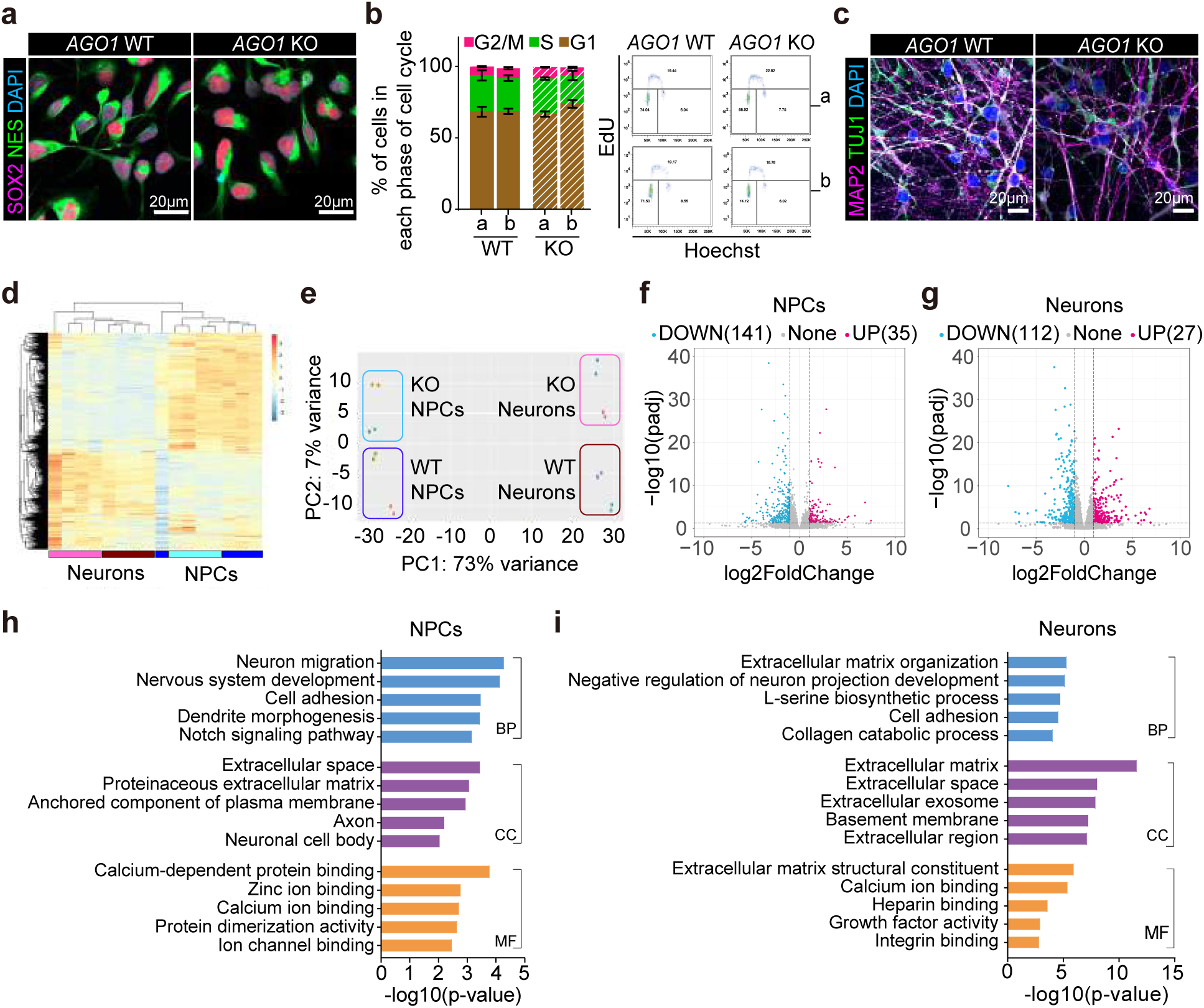
Comparative gene expression profiles of *AGO1* WT and KO hESC-derived NPCs and neurons. a. Representative confocal images from a single plane of NPCs stained for the NPC markers SOX2 (magenta) and NESTIN (NES; green), with DAPI (blue) indicating nuclei. Scale bar = 20 μm. b. Cell cycle analysis with EdU staining in *AGO1* WT and KO NPCs (n=3, mean ± SEM, two-way repeated measures ANOVA, *p*=0.9835). c. Representative confocal images from a single plane of 4-week differentiated neurons stained for the neuronal markers MAP2 (magenta) and TUJ1 (green) with DAPI (blue). Scale bar = 20 μm. (d, e) Heat map (d) and PCA plot (e) of RNA-seq results for *AGO1* WT and KO NPCs, as well as 4-week differentiated neurons. (f, g) Volcano plots of RNA-seq results for NPCs (f) and 4-week differentiated neurons (g). In NPCs, 176 DEGs (35 up, 141 down) and in neurons, 139 DEGs (27 up, 112 down) were identified based on criteria of |log2Foldchange|>1, adjusted p-value<0.05 (FDR<0.05), and WT_TPM_>0.5. (h, i) Gene ontology (GO) analysis of DEGs from NPCs (h) and 4-week differentiated neurons (i) using the DAVID database version 6.8. The analysis covers biological processes (BP), cellular components (CC), and molecular functions (MF).

Thus, to further explore the impact of *AGO1* KO in neuronal development, we performed total RNA sequencing (RNA-seq) using RNAs extracted from NPCs and 4-week differentiated neurons. The sequencing results, visualized through heatmaps and Principal Component Analysis (PCA) plots, showed clear group classifications (Fig. 3d,e). In our analysis of differentially expressed genes (DEGs) in *AGO1* KO cells, we identified 176 DEGs in NPCs and 139 DEGs in 4-week differentiated neurons (|log2FC|>1, padj<0.05) (Fig. 3f,g). Gene ontology (GO) analysis of these DEGs indicated that AGO1 regulates expression of genes crucial for cell adhesion and neuronal development in both NPCs and neurons (Fig. 3h,i). Cell adhesion, the process by which individual cells attach to each other and to the extracellular matrix, is essential in various aspects of neuronal development^37–39^.

We validated impaired cell adhesion in NPCs by focusing on NPCs within neural rosettes, as there were no observable issues when culturing monolayer NPCs on poly-L-ornithine (PLO)/laminin (Lam)-coated plates. NPCs cultured in a monolayer originate from the dissociation of neural rosettes into single cells, a process during which they lose the structural integrity typically maintained by intercellular adhesions (Extended Data Fig. 2a). Within these neural rosettes, NPCs are aligned with polarity around a centrally located lumen (Fig. 4a). To assess the structural integrity of NPCs in neural rosettes, we stained them with ZO-1 and N-cadherin antibodies, which mark tight and adherens junctions, respectively, at the apical junctions of NPCs facing the lumen^41^. Immunocytochemistry results from ZO-1 staining revealed that *AGO1* KO leads to morphological defects in the lumen of neural rosettes (Fig. 4b). The circularity of the lumen, calculated by the ratio of the minor axis to the major axis, was reduced in *AGO1* KO neural rosettes, indicating a more elongated oval shape (Fig. 4c). Similarly, the ventricle-like structures in brain organoids, stained with N-cadherin, also exhibited abnormal morphology in *AGO1* KO (Extended Data Fig. 3a).

**Fig. 4:**
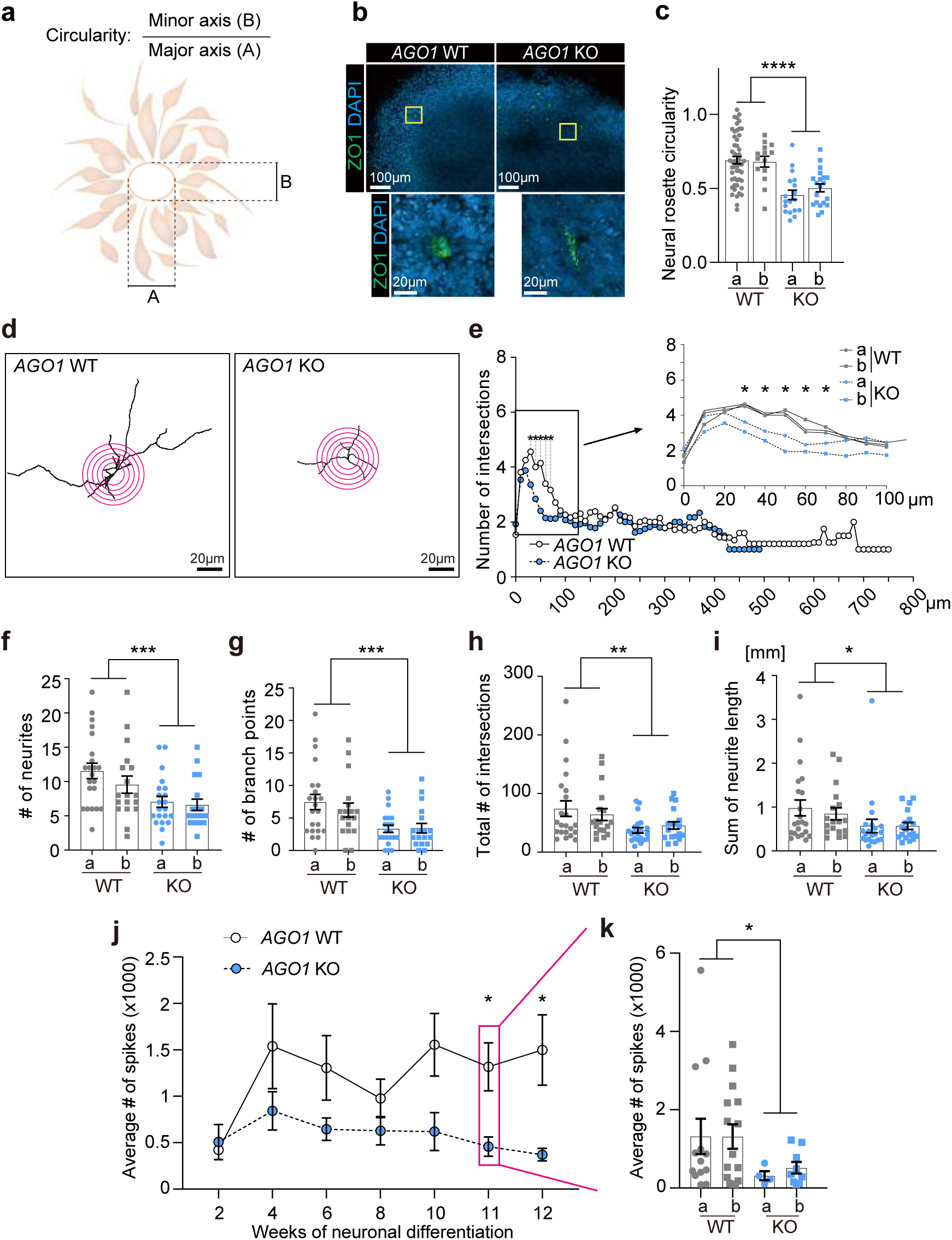
Cellular phenotypes of neural cells caused by AGO1 KO. a. Schematic diagram illustrating the definition of neural rosette circularity b. Representative confocal images from a single plane of neural rosettes used to analyze neural rosette circularity shown in (c). Circularity was analyzed by staining with ZO-1 (green), a tight junction marker in neural rosettes. DAPI is shown in blue. Scale bar = 50 μm (low magnification) and 20 μm (high magnification) .c. Quantification of circularity in *AGO1* WT and KO neural rosettes, where each dot represents a result from a single rosette structure. The following numbers of neural rosettes from 4 biological replicates were analyzed: n=37 WT_a, n=13 WT_b, n=31 KO_a, n=12 KO_b (mean ± SEM, unpaired ordinary one-way ANOVA with Tukey’s multiple comparisons test, \*\*\*\**p*<0.0001). d. Representative images of *AGO1* WT and KO neurons differentiated for 4 weeks, used for Sholl analysis, as shown in (e). Scale bar = 20 μm. e. Sholl analysis of *AGO1* WT and KO neurons Each dot represents the average number of intersections at a given distance from the soma of all analyzed neurons. Neurons from 3 biological replicates were analyzed. (n=22 WT_a, n=18 WT_b, n=21 KO_a, n=18 KO_b, mean ± SEM, multiple unpaired t-test, \**p*=0.0212 at 30 μm, \**p*=0.021 at 40 μm, \**p*=0.003 at 50 μm, \**p*=0.0086 at 60 μm, \**p*=0.0262 at 70 μm). (f-i) Morphology analysis results of *AGO1* WT and KO neurons, using the same neurons from the Sholl analysis in (e). Each dot in the graphs represents the value of a parameter for a single neuron (mean ± SEM, an unpaired ordinary one-way ANOVA with Tukey’s multiple comparisons test). f. Number of neurites (\*\**p*=0.0024). g. Number of branch points (\*\**p*=0.0031). h. Total number of intersections (\*\**p*=0.0023). i. Sum of total neurite length (\**p*=0.0402) j, k. Average number of spikes in *AGO1* WT and KO neurons analyzed by MEA. The spike number detected in a well was calculated by averaging the spike counts from all active electrodes within that well. Experiments were performed in 3 independent biological replicates. j. Changes in the average number of spikes during neuronal differentiation. Spike numbers from multiple wells were averaged from 3 biological replicates at each time point of neuronal differentiation. The number of wells varies by time point. (n=9-19 WT_a, n=2-18 WT_b, n=3-12 KO_a, n=6-19 KO_b, mean ± SEM, multiple unpaired t-test, \**p*=0.0327 at 11-week, \**p*=0.0415 at 12-week). k. Average number of spikes in 11-week differentiated neurons. Each dot represents the number of spikes detected in a well. The total number of wells analyzed is as follows. (n=13 WT_a, n=15 WT_b, n=4 KO_a, n=9 KO_b) (mean ± SEM, unpaired Student’s t-test, \**p*=0.0359).

The defects in neuronal development resulting from *AGO1* KO were confirmed via morphological analysis of neurons differentiated for 4 weeks. Using Sholl analysis, which involved counting neurite intersections at 10-μm intervals from the soma (Fig. 4d), we noted a significant reduction in neurite outgrowth and interactions in *AGO1* KO neurons compared to the control (Fig. 4e). Furthermore, a decrease in all parameters, including the total number of neurites, branch points, interactions, and total neurite length, was observed in *AGO1* KO neurons, indicating reduced neurite complexity (Fig. 4f-i). These findings demonstrate that AGO1 contributes to neuronal development by influencing neurite outgrowth and maturation. The effects of *AGO1* KO on the developmental deficits of neurons were further supported by electrophysiological properties observed in *AGO1* KO neurons. Multielectrode array (MEA) analysis revealed a decrease in spike numbers at all time points during neurogenesis (Fig. 4j, k). Similar patterns were observed in mean firing rates and burst numbers, both of which were reduced in *AGO1* KO neurons (Extended Data Fig. 3b-e). These electrophysiological changes were corroborated by raster plots from 11-week differentiated neurons (Extended Data Fig. 3f).

The cellular phenotypes of *AGO1* KO neural cells produced by the 2D culture method align with those observed in *AGO1* KO organoids (Fig. 2d-g, 4a-c, and Extended Data Fig. 1d-f,3a). The disorganized ventricle-like structures in *AGO1* KO brain organoids resemble the abnormal structure of neural rosettes produced by the 2D culture method, likely due to impaired cell adhesion. The disruption of the junctional complex leads to diminished polarity in NPCs, resulting in a deficit of neuronal development. Similarly, in the organoids, delayed neuronal development is evidenced by thinner cortical layers, which are comparable to those observed in 2D cultured neurons with short neurites and reduced neuronal activity.

### AGO1 represses LIN28A expression at the transcriptional level in NPCs

Next, we investigated the molecular mechanisms by which AGO1 influences cell adhesion and neuronal development. Initially, we assessed AGO1 protein expression levels across different stages of neuronal development, finding the highest levels in NPCs via Western blot analysis (Fig. 5a). This finding suggests that the effects of *AGO1* KO may be most pronounced in NPCs, compared to earlier or later developmental stages. Additionally, considering that AGO1 performs distinct roles in the cytoplasm and nucleus, regulating gene expression post-transcriptionally in the cytoplasm and transcriptionally in the nucleus, we explored its primary subcellular localization within neural cells. The results of nuclear-cytoplasmic fractionation showed that a substantial amount of AGO1 protein is localized within the nucleus (Fig. 5b). Based on these findings, we hypothesized that AGO1 likely plays significant roles in the nucleus of NPCs, particularly in regulating cell adhesion and neuronal development.

**Fig. 5:**
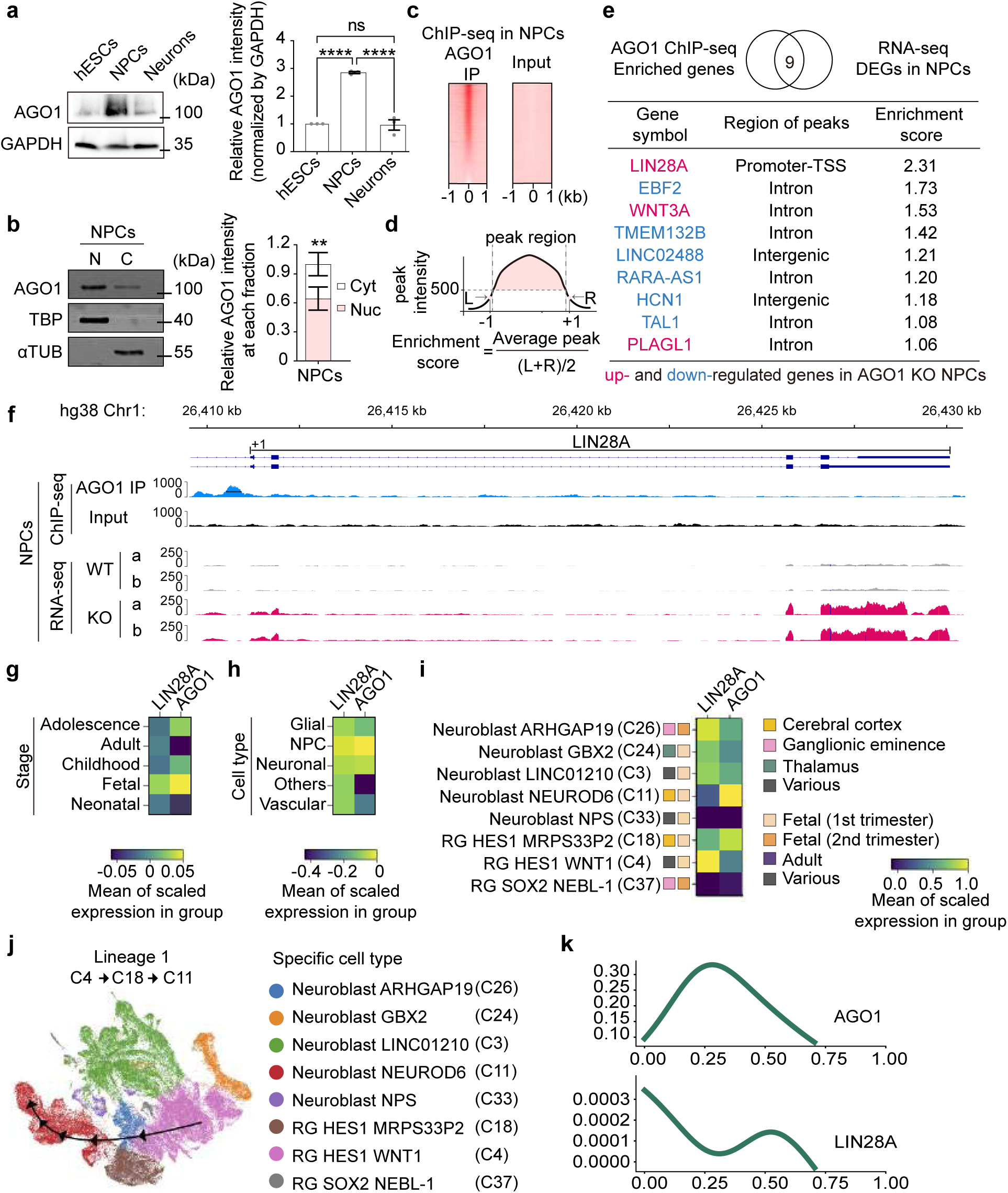
Transcriptional repression of LIN28A by nuclear AGO1 in NPCs. a. Representative Western Blot images showing AGO1 expression levels during neuronal differentiation. GAPDH (∼35 kDa) was used as a loading control. Quantified AGO1 band intensities from 3 replicates were normalized to GAPDH (mean ± SEM, unpaired ordinary one-way ANOVA with Tukey’s multiple comparisons test; hESCs vs. NPCs \*\*\*\**p*<0.0001, hESCs vs. neurons *p*=0.9651, NPCs vs. neurons \*\*\*\**p*<0.0001). b. Representative Western Blot images showing the subcellular localization of AGO1 in NPCs. TATA binding protein (TBP) and αTUB were used as controls for nuclear and cytoplasmic fractions, respectively. Quantified AGO1 band intensities from 3 replicates were normalized to their respective fractional control (mean ± SEM, unpaired one-way ANOVA with Sidak’s multiple comparisons test; \*\**p*=0.0034). c. Heatmap of AGO1 ChIP-seq results in NPCs. d. Diagram depicting the calculation of enrichment score. The enrichment score is calculated by dividing the average intensity (INT) of the peak region by the average INT of the flanking regions located-1 bp(L) and at +1 bp (R) from the peak boundaries. The average INT of the peak region is determined by dividing the total INT of the peak region by the length of the peak. e. List of genes enriched in AGO1 ChIP-seq and differentially expressed in *AGO1* KO NPCs. The gene closest to the peaks with an enrichment score was identified as the enriched gene from ChIP-seq results. f. Visualization of AGO1 ChIP-seq peaks and RNA-seq reads on the LIN28A genomic region using Integrative Genomics Viewer. (g-k) Expression patterns of AGO1 and LIN28A during human brain development. Single-cell RNA sequencing data from various stages of human brain development were analyzed using the Single Cell Brain Atlas. g. Heatmap displaying AGO1 and LIN28A expression levels across different developmental stages of the human brain. h. Heatmap displaying AGO1 and LIN28A expression levels across classified cell types of the human brain. i. Heatmap of AGO1 and LIN28A expression levels in NPC clusters from different brain regions and developmental stages, with each cluster categorized by a specific marker. j. A cell differentiation lineage (Lineage 1) showing anti-correlated expression patterns between AGO1 and LIN28A. k. Expression patterns of AGO1 and LIN28A along the Lineage 1 trajectory pseudo-time.

Nuclear AGO1 regulates gene expression by associating with chromatin^42,43^. To explore the genomic locations of AGO1 binding in NPCs, we conducted chromatin immunoprecipitation sequencing (ChIP-seq) and calculated enrichment scores (Fig. 5c,d, and Extended Data Fig.4a). By comparing genes located within 1kb of AGO1 binding sites (from ChIP-seq data) with DEGs (from RNA-seq data), we identified 9 genes potentially impacted by AGO1 binding (Fig. 5e). Among these, LIN28A was particularly noteworthy, where AGO1 binds at the promoter-TSS (transcription start site) with the highest enrichment score. LIN28A also emerged as one of the top 5 upregulated genes in *AGO1* KO NPCs (Extended Data Fig. 4b). We validated the increase in LIN28A expression at both the mRNA and protein levels in *AGO1* KO NPCs (Extended Data Fig. 4c,d). To exclude the possibility of post-transcriptional regulation of LIN28A expression by AGO1, we blocked transcription in *AGO1* WT and KO NPCs using actinomycin D (Act. D) and compared mRNA decay rates (Extended Data Fig. 4e). The half-life of LIN28A mRNA was similar between both groups. These data indicate that AGO1 binds to the promoter-TSS region of LIN28A and suppresses the expression of LIN28A in NPCs (Fig. 5f).

The single cell atlas of the human brain revealed that both AGO1 and LIN28A are highly expressed in the fetal brain and within NPCs (Fig. 5g,h). A more detailed analysis of the expression patterns of AGO1 and LIN28A was conducted using 8 NPC clusters, which we categorized based on cell types, brain regions, and developmental stage. Notably, AGO1 presents the highest expression levels in neuroblasts in the cerebral cortex during the first trimester (Cluster 11, C11), whereas LIN28A shows peak expression in radial glia (RG) across various regions of the brain during the same period (C4) (Fig. 5i). Cell trajectory analyses indicated that C11 neuroblasts are differentiated from C4 RG through C18 RG, an RG cell type mainly located in the cerebral cortex. This process was designated as Lineage 1 among 4 identified cell differentiation lineages (Fig. 5j and Extended Data Fig. 4f). Intriguingly, AGO1 and LIN28A displayed clear anti-correlative expression patterns in Lineage 1 among 4 lineages, specifically at the early stages of differentiation pseudo-time (Fig. 5k and Extended Data Fig.4g). The expression patterns of AGO1 and LIN28A, acquired from the single cell brain atlas, support our finding that AGO1 suppresses the transcription of *LIN28A* in hESC-derived NPCs, which significantly resembles the transcriptional profiles of the first trimester of the human cortex (Extended Data Fig.4h).

### LIN28A suppresses REELIN at the post-transcriptional level in NPCs

LIN28A, an RBP, inhibits translation by binding to mRNAs being translated in the endoplasmic reticulum (ER)^44^. Given this function, we presumed that elevated LIN28A in *AGO1* KO NPCs might suppress the translation of ER-associated mRNAs, thereby reducing mRNA stability as these stalled mRNAs become more susceptible to degradation^45^. To verify this hypothesis, we analyzed DEGs for features indicative of ER translation and discovered that most DEGs encode proteins undergoing N-linked glycosylation, a co-translational modification within the ER (Fig. 6a). As expected, most genes (53 of 64) involved in N-linked glycosylation were downregulated in *AGO1* KO NPCs with increased LIN28A levels (Supplementary Table 1).

**Fig. 6:**
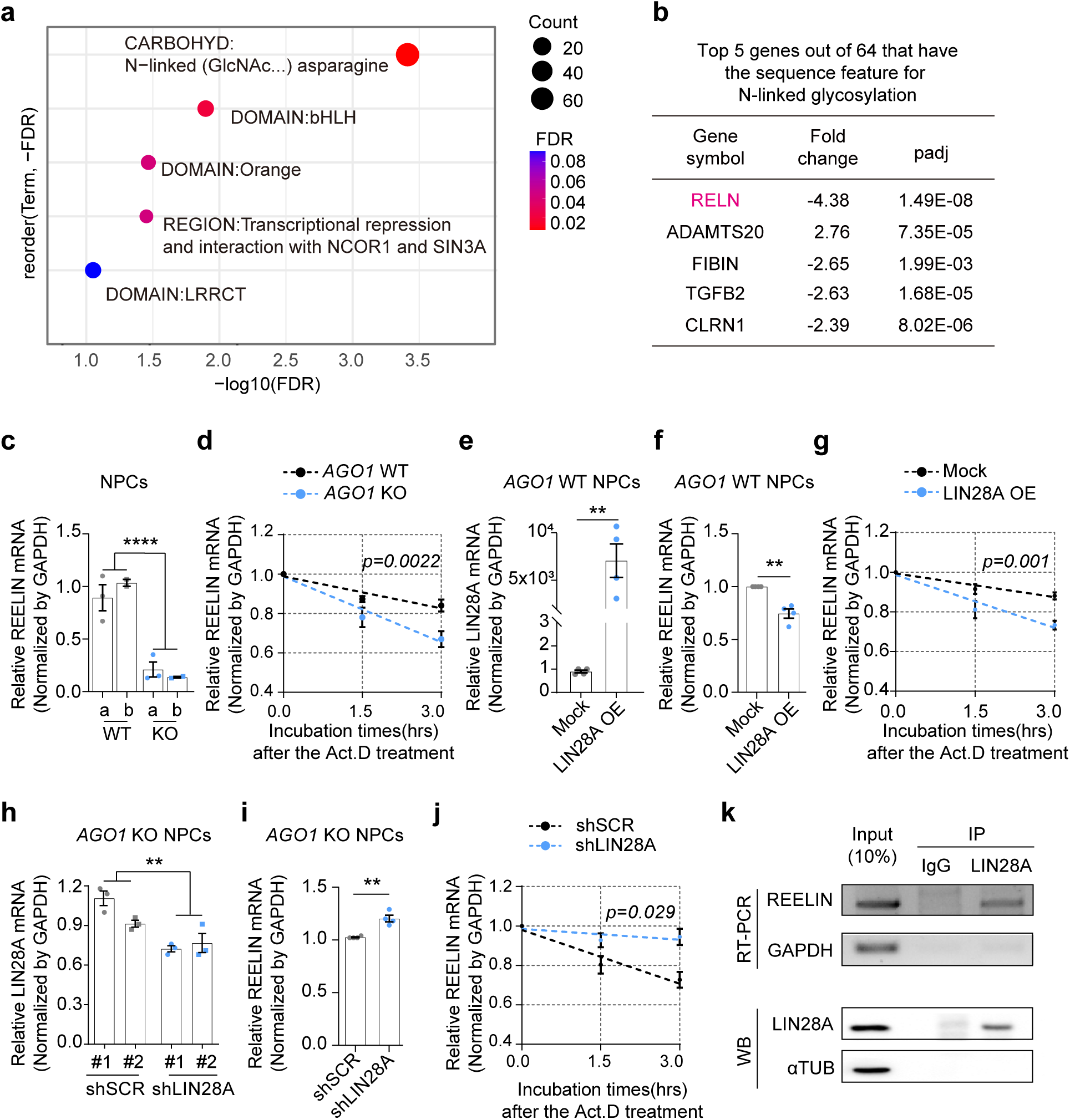
LIN28A-mediated reduction in REELIN mRNA stability in NPCs. a. UniProt sequence features of DEGs identified in *AGO1* KO NPCs. b. Top 5 genes exhibiting the largest fold changes among DEGs with N-linked glycosylation feature. c. Quantitative PCR (qPCR) of REELIN expression in *AGO1* WT and KO NPCs. (n=3 WT_a, n=2 WT_b, n=3 KO_a, n=2 KO_b, mean ± SEM, unpaired ordinary one-way ANOVA with Sidak’s multiple comparisons test, \*\*\*\**p*<0.0001). d. qPCR measurement of REELIN mRNA decay rate in *AGO1* WT and KO NPCs following transcription inhibition with Actinomycin D (Act. D). The analysis included results from 4 biological replicates. (n=4 WT: n=2 WT_a, n=2 WT_b; n=4 KO: n=2 KO_a, n=2 KO_b, mean ± SEM, \*\**p*=0.0022 in simple linear regression between the two samples). e. qPCR analysis of LIN28A expression in WT NPCs to confirm overexpression following LIN28A transfection (n=4 WT: n=2 WT_a, n=2 WT_b, mean ± SEM, unpaired Student’s t-test, \*\**p*=0.0071) f. qPCR of REELIN expression in WT NPCs following overexpression of LIN28A compared to control. (n=4 WT: n=2 WT_a, n=2 WT_b, mean ± SEM, unpaired Student’s t-test, \*\**p*=0.0037). g. qPCR measurement of REELIN mRNA decay rate in LIN28A-overexpressing and control NPCs following Act. D treatment (n=6 WT: n=3 WT_a, n=3 WT_b, mean ± SEM, \*\**p*=0.001 in simple linear regression between the 2 samples). h. qPCR analysis to confirm LIN28A knockdown in *AGO1* KO_a NPCs after lentiviral shRNA infection. Two different shRNAs targeting LIN28A mRNAs (shLIN28A) and 2 different scrambled shRNAs (shSCR) were utilized (n=3, mean ± SEM, unpaired ordinary one-way ANOVA with Tukey’s multiple comparisons test, \*\**p*=0.0021). i. qPCR of REELIN expression in *AGO1* KO_a NPCs following LIN28A knockdown. Results were obtained from biological duplicates using 2 shLIN28As and 2 shSCRs. (mean ± SEM, \*\**p*=0.0015, unpaired Student’s t-test). j. qPCR measurement of REELIN RNA decay rate in *AGO1* KO_a NPCs with LIN28A knockdown following Act. D treatment (n=3, mean ± SEM, \**p*=0.029 in simple linear regression between the 2 samples). k. Reverse transcription PCR (RT-PCR) of REELIN mRNA associated with LIN28A in NPCs following RNA immunoprecipitation (IP) using anti-LIN28A antibody. To confirm the specificity of the antibody, IgG was employed as a control. Expression levels of REELIN were quantified from total RNAs extracted from 10% of the total lysates used for RNA IP, referred to as input. GAPDH was used as a control.

It is noteworthy that REELIN emerged as the most significantly downregulated gene, displaying the highest fold change, not only among genes associated with N-linked glycosylation but also across all DEGs analyzed (Fig. 6b). REELIN, a glycoprotein, plays a critical role in neuronal development by regulating neuronal migration, dendritic growth, and synaptic functions^46–48^. We confirmed a decrease in REELIN mRNA levels in *AGO1* KO NPCs and further verified that this decrease is a result of reduced RNA stability at the post-transcriptional level (Fig. 6c). By measuring the half-life of REELIN mRNA, we observed that it decayed more rapidly in *AGO1* KO NPCs compared to *AGO1* WT NPCs (Fig. 6d).

To determine if the rapid REELIN mRNA decay in *AGO1* KO NPCs is due to enhanced LIN28A activity, we manipulated LIN28A expression levels in both *AGO1* WT and KO NPCs (Fig. 6e-j). Overexpressing LIN28A in *AGO1* WT NPCs led to reduced REELIN mRNA levels (Fig. 6f). This reduction in REELIN mRNA level in LIN28A-overexpressing NPCs was due to a decrease of RNA half-life (Fig. 6g). Conversely, suppressing enhanced LIN28A in *AGO1* KO NPCs increased REELIN mRNA levels (Fig. 6i), and LIN28A knockdown in these cells extended the half-life of REELIN mRNA (Fig. 6j). These results indicate that the decrease in REELIN mRNA stability in *AGO1* KO NPCs is dependent on LIN28A expression levels.

To examine whether LIN28A directly binds to REELIN mRNA and reduces its stability, we conducted LIN28A RNA immunoprecipitation and performed reverse transcription-PCR using RNAs extracted from the ribonucleoprotein complex. The results showed that LIN28A specifically binds to REELIN mRNA, but not to GAPDH mRNA, which served as a control (Fig. 6k). Our data showed that increased LIN28A in *AGO1* KO NPCs suppresses REELIN, translated in the ER, by binding to their mRNAs.

### Reversal of *AGO1* KO-induced NPC phenotypes through restoration of LIN28A and REELIN expression

Finally, we investigated whether LIN28A and REELIN are the key downstream genes of AGO1 responsible for the observed phenotype in *AGO1* KO NPCs. To this end, we restored LIN28A and REELIN expression in *AGO1* KO NPCs and analyzed the circularity of the lumen in neural rosettes (Fig. 7). LIN28A, which is upregulated in *AGO1* KO NPCs, was downregulated using shRNA targeting LIN28A (Fig.7a,b). Conversely, REELIN levels were restored by treating the *AGO1* KO neural rosettes with recombinant REELIN protein (Fig. 7c,d). Reducing LIN28A expression via shRNA or treating with recombinant REELIN protein corrected the elongated oval shape of the lumen in neural rosettes, resulting from *AGO1* KO, to a round shape comparable to that observed in WT neural rosettes. These findings affirm that the AGO1-LIN28A-REELIN regulatory axis is critical for maintaining NPC polarity, which in turn influences neuronal development efficiency (Extended Data Fig.5).

**Fig. 7:**
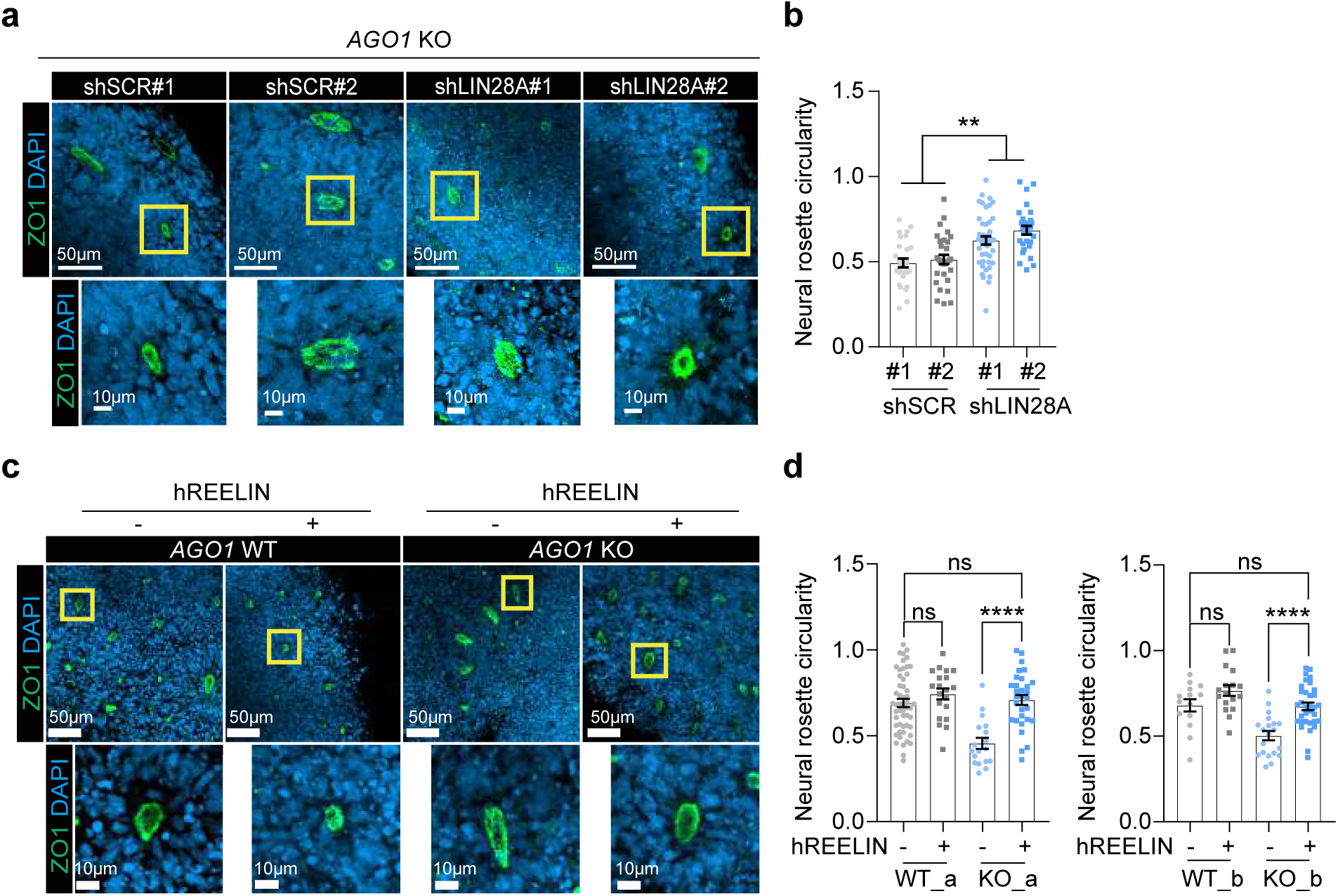
Restoration of NPC phenotypes caused by AGO1 KO via REELIN and LIN28A. (a, b) Neural rosette circularity in *AGO1* KO neural rosettes following LIN28A knockdown. a. Representative confocal images from a single plane of neural rosettes stained with the tight junction marker ZO-1 (green) and DAPI (blue), quantified in (b). Scale bar = 50 µm (low magnification) and 10 µm (high magnification). b. Quantification of neural rosette circularity, with each dot representing the result from a single rosette. The following numbers of neural rosettes from 3 biological replicates were analyzed: n=27 shSCR#1, n=30 shSCR#2, n=44 shLIN28A#1, n=28 shLIN28A#2 (mean ± SEM, unpaired ordinary one-way ANOVA with Tukey’s multiple comparisons test, \*\**p*=0.0022) (c, d) Neural rosette circularity in *AGO1* WT and KO rosettes, treated with or without human recombinant REELIN. c. Representative confocal images from a single plane of neural rosettes stained with the tight junction marker ZO-1 (green) and DAPI (blue), quantified in (d). Scale bar = 50 µm (low magnification) and 10 µm (high magnification). d. Quantification of neural rosette circularity, with each dot representing the result from a single rosette. The following numbers of neural rosettes from 3 biological replicates were analyzed: WT_a (n=52 no REELIN, n=20 with REELIN), KO_a (n=18 no REELIN, n=30 with REELIN), WT_b (n=14 no REELIN, n=17 with REELIN), KO_b (n=20 no REELIN, n=33 with REELIN) (mean ± SEM, unpaired ordinary one-way ANOVA with Tukey’s multiple comparisons test; WT_a vs. WT_a +REELIN *p*=0.6232, KO_a vs. KO_a +REELIN \*\*\*\**p*<0.0001, WT_a vs. KO_a +REELIN *p*=0.9806, WT_b vs. WT_b +REELIN *p*=0.2463, KO_b vs. KO_b +REELIN \*\*\*\**p*<0.0001, WT_b vs. KO_b +REELIN *p*=0.9990).

## Discussion

In this study, we uncovered the biological roles of AGO1 in organizing cortical architecture and influencing social behaviors. Conditional KO of *Ago1* in the brain leads to less sociable behavior in animals. Furthermore, depletion of *AGO1* disrupts the structure of ventricle-like formations in human forebrain organoids and delays neuronal development. The disruption of cortical structures, a characteristic feature observed in the brains of ASD patients^49^ and in ASD animal models developed under conditions of maternal immune activation^50^, corresponds to our findings of abnormal cortex development. Additionally, reduced electrophysiological properties in *AGO1* KO neurons, similar to those reported in neurons derived from ASD patient iPSCs, further underscore the critical role of AGO1 in neuronal function^51^.

Our findings underscore the unique functional characteristics of AGO1 in brain development and associated diseases, distinguishable from other proteins in the AGO family. While all mammalian AGO proteins (AGO1-4) load various small noncoding RNAs and generally target similar mRNAs, suggesting a potential functional redundancy within the RNA-induced silencing complex (RISC)^52–54^, *AGO1* KO reveals unique phenotypic consequences not compensated for by AGO2-4. These results emphasize the critical and non-redundant roles of AGO1 within its protein family, highlighting its unique contributions to brain structure and function. The specific impact of AGO1 among the 4 AGO proteins on brain development may originate from its nuclear localization. We discovered that AGO1 is predominantly localized in the nuclei of NPCs and is associated with chromatin (Fig. 5b). AGO1 displays distinct nuclear distribution patterns compared to AGO2^55^ and can regulate gene expression with or without small RNAs, unlike AGO2, which requires these RNAs for nuclear gene regulation^27,42,56–63^. During human brain development, the expression pattern of AGO1 shows a high correlation with genes responsible for chromatin remodeling^19^, further supporting the significance of the nuclear function of AGO1 in brain development and ASD.

In human NPCs, AGO1 targets the promoter-TSS region of LIN28A, resulting in transcriptional suppression (Fig. 5e). LIN28A, an RBP that is highly expressed in NPCs during nervous system development^64,65^, regulates gene expression in versatile ways. Depending on the cellular context, LIN28A can either suppress or enhance translation of mRNAs^66^. The regulatory actions of LIN28A on mRNA translation can function independently or in conjunction with microRNAs, such as the let-7 family, whose production is also regulated by LIN28A^67–70^. Additionally, LIN28A can induce the decay of mRNAs by binding to them directly^66^. Our RNA-seq results reveal that most DEGs in *AGO1*-deficient NPCs are likely translated through the ER (Fig. 6a). Although in mouse ESCs, LIN28A-mediated translational suppression of ER-associated mRNAs does not alter mRNA expression levels^44^, our results indicate a significant reduction in the expression of mRNAs translated through the ER in *AGO1* KO NPCs, where LIN28A levels are elevated. Our data also show a notable decrease in most DEGs in *AGO1*-deficient NPCs (Fig. 3f), indicating the possibility that an increase in LIN28A might lead to the rapid decay of a large number of mRNAs. We demonstrated LIN28A-mediated mRNA decay at the post-transcriptional level using REELIN, the most downregulated mRNA in *AGO1*-deficient NPCs (Fig. 6d). Although REELIN is a major regulatory molecule that was the focus of our study, further investigation into whether LIN28A directly binds to and regulates the stability of other mRNAs in NPCs would be worthwhile.

We propose that REELIN is the key effector molecule regulated by AGO1 via LIN28A in human NPCs. The application of recombinant REELIN successfully restored the characteristic narrow and elongated lumen of *AGO1* KO neural rosettes, indicative of reduced patterning^71^ (Fig. 7c,d). REELIN, crucial in governing the cytoarchitecture patterning of the developing brain, is also reported as a high-confidence ASD risk factor. Beyond genetic variations, REELIN expression levels are consistently annotated as DEGs in numerous ASD studies^72^. Moreover, impaired REELIN signaling is a recurring observation in patients with ASD^73^, underlining the pivotal role of REELIN in ASD pathophysiology. Our findings align with these previous observations, demonstrating that alterations in REELIN expression can occur independently of genetic variations in the *RELN* gene. This independence of genetic variation highlights the potential of targeting REELIN pathways for therapeutic interventions in ASD.

Patients with ASD and other NDDs who carry genetic variations in the AGO1 gene display heterozygous mutations in one allele. In our study, although we utilized *Ago1* KO models, we found significantly relevant phenotypes of ASD. However, it remains uncertain whether these variations lead to dominant negative effects, loss-of-function, or gain-of-function impacts^74^, as the effects can vary depending on the specific mutations. Recently, AGO1 variations identified in human NDD patients were introduced into ALG-1 in *Caenorhabditis elegans*^24^, the homologous protein of human AGO1, revealing that some mutations result in more severe developmental phenotypes than null. These ALG-1 mutations result in heterochronic developmental phenotypes, varying widely among different mutations. Moving forward, it would be interesting to introduce each variation of the *AGO1* gene into mouse models and human neural cell models to determine their specific impacts on brain development. Identifying how these *AGO1* variations affect brain development differently in each individual will be crucial for designing personalized medicine approaches.

## Supplementary Figure legends

**Extended Data Fig. 1: Phenotypes of *AGO1* KO forebrain organoids**

a. Growth curve of forebrain organoids throughout the differentiation period.

b. Representative bright-field images of forebrain organoids during differentiation. Scale bar = 500 μm.

c. Representative confocal images from a single plane of a day 50 differentiated *AGO1* WT organoid, with DAPI (blue). A substructure within the yellow box is defined as a ventricle-like structure. Scale bar = 500 μm.

d. Confocal images of a ventricle-like structure stained with cortical layer-specific markers: PAX6 (magenta) for the ventricular zone (VZ) and CTIP2 (white) for the cortical plate (CP). The layer resembling the subventricular zone (SVZ) shows co-expression of PAX6 and CTIP2. Scale bar = 500 μm.

(e, f) Statistical analysis of quantified relative layer thickness from specific marker staining in forebrain organoids, as shown in Fig. 2g, which combined results from both cell lines. Results for individual cell lines ‘a’ and ‘b’ are now presented separately. Data are shown as mean ± SEM and analyzed using an unpaired two-way ANOVA with Sidak’s multiple comparisons test.

e. n=11 WT_a, n=12 KO_a, VZ-like layer, \**p*=0.0170; SVZ-like layer, \**p*=0.0446; CP-like layer, *p*=0.9774.

f. n=3 WT_b, n=5 KO_b, VZ-like layer, \*\*\**p*=0.0003; SVZ-like layer, \**p*=0.0228; CP-like layer, *p*=0.2027.

**Extended Data Fig. 2: Differentiation of *AGO1* WT and KO hESCs into NPCs and neurons**

a. Schematic of the protocol for differentiating hESCs into NPCs and neurons.

(b, c) Proportion of cells expressing cell type-specific markers in NPCs and neurons differentiated from *AGO1* WT and KO hESCs, across three biological replicates. Results are presented as mean ± SEM, analyzed using an unpaired ordinary one-way ANOVA with Tukey’s multiple comparisons test.

b. Proportion of SOX2 and NESTIN (NES) double-positive cells in NPCs. Total number of cells counted using DAPI (n=382 WT_a, n=585 WT_b, n=625 KO_a, n=663 KO_b, unpaired ordinary one-way ANOVA with Sidak’s multiple comparisons test, *p*=0.2304).

c. Proportion of TUJ1 and MAP2 double-positive cells in neurons differentiated for a week. Total number of cells counted using DAPI (n=287 WT_a, n=178 WT_b, n=260 KO_a, n=211 KO_b, unpaired ordinary one-way ANOVA with Sidak’s multiple comparisons test, *p*=0.3169).

**Extended Data Fig. 3: Phenotype analysis of neural cells resulting from *AGO1* KO**

a. Representative confocal images from a single plane of day 50 forebrain organoids. N-cadherin (NCAD), a neuroepithelial tube marker, is shown in magenta, and DAPI in white. Scale bar = 20 μm. (b-e) Additional electrophysiological features of *AGO1* WT and KO neurons analyzed using a MEA, based on the same dataset as in Fig. 4j-k.

b. Mean firing rates during neuronal development. (mean ± SEM, multiple unpaired t-test, \**p*=0.0327 at 11-week, \**p*=0.0415 at 12-week).

c. Mean firing rates at 11-week differentiated neurons. (mean ± SEM, unpaired Student’s t-test, \**p*=0.0359).

d. Average number of bursts during neuronal development. (mean ± SEM, multiple unpaired t-test, \*\**p*=0.0073 at 10-week, \**p*=0.0044 at 11-week, \**p*=0.0054 at 12-week).

e. Average number of bursts at 11-week differentiated neurons. (mean ± SEM, unpaired Student’s t-test, \*\**p*=0.0048)

f. Representative raster plots of 11-week differentiated neurons. Black lines indicate spikes, blue lines indicate bursts, and purple boxes represent network bursts.

**Extended Data Fig. 4: Roles of AGO1 suppressing LIN28A in NPCs**

a. Annotated genomic regions and the number of peaks at the given regions identified from AGO1 ChIP-seq analysis in NPCs.

b. Top 5 up-regulated and down-regulated DEGs based on fold change in AGO1 KO NPCs.

c. qPCR of LIN28A expression in *AGO1* WT and KO NPCs. (n=4, mean ± SEM, unpaired ordinary one-way ANOVA with Sidak’s multiple comparisons test, \*\**p*=0.0013).

d. Representative Western Blot images showing LIN28A expression level in NPCs. αTUB was used as a loading control. Quantified LIN28A band intensities from 3 replicates were normalized to αTUB. (mean ± SEM, unpaired ordinary one-way ANOVA with Sidak’s multiple comparisons test, \*\**p*=0.0021).

e. qPCR measurement of LIN28A mRNA decay rate in *AGO1* WT and KO NPCs following Act. D treatment. (n=4 WT: n=2 WT_a, n=2 WT_b; n=4 KO: n=2 KO_a, n=2 KO_b, mean ± SEM, *p*=0.7856 in simple linear regression between the 2 samples).

f. Additional cell differentiation lineages (Lineages 2-4) used to analyze the expression correlation between AGO1 and LIN28A, as shown in Fig. 5k.

g. Expression patterns of AGO1 and LIN28A along the trajectory time of each lineage.

h. Comparison of RNA expression profiles between hESC-derived NPCs and Neurons versus the human brain across developmental stages

**Extended Data Fig. 5: Summary of this study illustrating AGO1-LIN28A-REELIN regulatory axis in the cortical development**

## Materials and methods

### Animals

The *Ago1^tm1a^*(knockout first allele) mouse, designated as C57BL/6N-*Ago1^tm1a(KOMP)Wtsi^*/Mmucd (RRID: MMRRC_046532-UCD), was obtained from the Mutant Mouse Resource and Research Center (MMRRC) at University of California at Davis, an NIH-funded strain repository. The *Ago1^tm1a^* was donated to the MMRRC by The KOMP Repository, University of California, Davis, originating from Ramiro Ramirez-Solis, CSD. The *Protamine-FLP* (flippase) mouse was obtained from the Laboratory Animal Resource and Research Center (LARRC) at Korea Research Institute of Bioscience and Biotechnology (LARRC ID: AZ00000694). The *Nestin-Cre* transgenic mouse was acquired from Jackson Laboratory (stock: 003771). The *Ago1^tm1a^*was crossed with *Protamine-FLP* to generate the *Ago1^tm1c^*(conditional allele). Further, *Ago1^tm1c^* was crossed with *Nestin-Cre* to generate *Ago1^tm1d^* (deletion allele), a conditional *Ago1* KO in the brain. The mouse housing facility maintains specific pathogen-free conditions, with a constant temperature of 23°C and a 12-hour light-dark cycle, where the lights are turned off at 19:00.

### Mouse behavior tests: open field test and direct interaction test

Prior to the behavioral assessments, mice were habituated in the test room for at least 30 minutes. The tests were conducted in the following order: open field test and then direct interaction test, all during the dark cycle and within the same chamber. For the open field test, mice were gently placed in the center of the chamber, and the total distance moved and the time spent in the center were recorded over a 15-minute period. Subsequently, the direct interaction test was conducted by introducing a sex-matched novel mouse. Both the frequency and duration of sniffing by the test mouse towards the novel mouse were recorded over a 10-minute period. All behavioral tests were performed using male mice of *Ago1* WT and cKO, aged 8-9 weeks, with sex-matched novel mice aged 6-7 weeks. All experimental procedures received approval from the KAIST Institutional Animal Care and Use Committee (IACUC) (KA2022-035).

### hESCs culture

All the experiments were performed using H9 (WA09) hESCs, obtained from WiCell. The hESCs were maintained at 37°C and 5% CO_2_ as previously described, cultured in DMEM F-12 (Gibco 12400024) supplemented with 20% of Knockout serum replacement (KSR; Gibco 10828028), 1% MEM non-essential amino acids (NEAA; Gibco 11140050), 0.1 mM 2-mercaptoethanol (2-ME; Sigma M3148), 1.2 mg/ml sodium bicarbonate (Sigma S5761), and 10 ng/ml FGF2 (R&D systems 4114-TC) on mouse embryonic fibroblasts (MEFs), inactivated with 10 μg/ml of mitomycin C (AG Scientific M-1108) for 2 hours. The hESCs were maintained with daily media changes and passaged every 6 days to expand cells. When single ESCs needed to be selected and expanded, they were dissociated with Accutase, plated onto Matrigel (BD 354230) coated plates, and cultured in mTeSR (STEMCELL Technologies ST85850) with FGF2 (R&D systems 4114-TC) and Y27632 (Selleckchem S1049). Once a single ESC was selected and had grown to form a colony, the ESCs were cultured again on MEFs. All studies were conducted using hESCs at passage numbers less than 65. The Institutional Review Board of KAIST has approved the protocols for using hESCs in accordance with the relevant ethical standards and regulations (KH2021-069).

### *AGO1* KO in hESCs

Single guide RNAs (sgRNAs), targeting sequences 5’-GTGGCTAGCCATTGTGAGC-3’ in the second constitutive exon and 5’-TGAGGGCTACTACCACCCGC-3’ in the third constitutive exon of the *AGO1* gene, were designed as sgRNA_a and sgRNA_b, respectively, and cloned into the pSpCas9(BB)-2A-GFP (pX458) vector, developed by Feng Zhang (Addgene 48138)^75^. hESCs were dissociated into single cells and transfected with these plasmids using a nucleofection kit (Lonza Bioscience, VPG-1005). Cells expressing GFP were isolated by FACS sorting and plated in 24-well plates to grow and form colonies. The genomic DNA was extracted from single colonies and amplified for genotyping by Sanger sequencing using the following primers: KO_a_Fwd 5’-CAGAAGCACTGAGCCAAGGTG-3’; KO_a_Rev 5’-AGCAGTTCTGCCAACACTAGTAGC-3’; KO_b_Fwd 5’-TGCCCAGGATGCCTCACAGG-3’; KO_b_Rev 5’-GTGCCAGGGCTGTAGGAATTAGAC-3’

### Neuronal differentiation of hESCs using 2D methods

To produce neural rosettes, hESC colonies were dissociated using 10 μg/ml collagenase IV (Gibco 17104019) in DMEM F-12 at 37°C and 5% CO_2_ for 1 hour until colonies detached from the dish. The hESCs were then transferred to non-adherent dishes and maintained in hESC culture media. The following day, the media were replaced with embryoid body (EB) media containing dual SMAD inhibitors [10 μM of SB-431242 (Cayman CAY-13031) and 0.1 μM of LDN193189 (Selleckchem S2618) in N2B27 media (DMEM F-12/glutamax (Gibco 10565042), supplemented with N2 (Gibco 17502048), and B27 (Gibco12587010)]. The floating cell culture was continued for 7-10 days to form EBs, which were then plated onto Matrigel-coated dishes with EB media with 1 μg/ml of Lam (Gibco 23017015) to facilitate attachment. After 3-5 days, neural rosettes became visible.

To generate NPCs, these neural rosettes were manually picked, dissociated with Accutase (Innovative Cell Technologies AT104) and plated onto PLO (Sigma P3655)/Lam-coated dishes in N2B27 media supplemented with 20 ng/ml of FGF2. NPCs were maintained at a high density (150k-250k cells/cm^2^) and split approximately 1:3 to 1:4 every 1-2 weeks using Accutase.

To differentiate NPCs into neurons, the cells were plated at low density (30k-40k cells/cm^2^) onto plates coated with PLO/Lam. The following day, culture media were changed by removing FGF2 and adding 1 μg/ml Lam, 20 ng/ml BDNF (PeproTech 450-02), 20 ng/ml GDNF (PeproTech, AF-450-10), 500 μg/ml of dibutyryl-cyclic AMP (Sigma D0627), and 200 nM ascorbic acid (Sigma A4034). During neuronal differentiation, half of the culture media was replaced with fresh media every 2 days.

### Forebrain organoid differentiation

Forebrain organoids were differentiated using a method previously described, with minor modifications^76^. hESCs, dissociated with collagenase IV, were transferred to non-adherent dishes and maintained in organoid media (i) containing 2 nM Dorsomorphin (STEMCELL Technologies 72102), 2 nM A-83 (STEMCELL Technologies 72022), 1% MEM NEAA, 0.1 mM 2-ME, 20% KSR, and 1% penicillin/streptomycin (PS) (Welgene LS202-02) in DMEM F-12/Glutamax. The following day (Day 1), the media were refreshed. On Days 3 and 4, half of the culture media was replaced with fresh media. On Days 5 and 6, half of the culture media was replaced with fresh organoid media (ii), including 1 nM CHIR-99021(STEMCELL Technologies 72052), 0.1 nM SB-431242, 1% MEM NEAA, 1% PS, and N2 supplement in DMEM F-12/Glutamax. On Day 7, 4-5 organoids were embedded in 30 µl Matrigel on parafilm and incubated at 37°C and 5% CO_2_ for 30 minutes. The Matrigel-coated organoids were then transferred into non-adherent dishes and cultured with organoid media (ii). Half of the culture media was replaced with fresh media every 2 days. On Day 14, mechanical dissociation of organoids from the Matrigel was performed. Thereafter, the organoids were cultured on a shaker in N2B27 media supplemented with 2-ME, 1% PS, and 2.5 ng/ml insulin (Sigma I9278) until Day 50.

### Cell cycle analysis

For cell cycle analysis, cells were prepared following manufacturer’s instructions (Invitrogen C10634). Briefly, *AGO1* WT and KO NPCs were treated with 10 µM EdU-containing cell culture media for 1 hour in an incubator (37 °C, 5% CO_2_). EdU-labeled cells were harvested and washed once with DPBS (Welgene, LB201-02) after the incubation, then pelleted down to remove the supernatant. The following sample preparation steps were performed in the dark at room temperature (RT) and washed with 1% BSA (Gendepot A0100-10) in DPBS. Pellet down step was executed at 1,440 g, 4 minutes at RT. The cell pellet was then dislodged and fixed with a Click-iT fixative for 15 minutes. After the fixation, cells were washed once and pelleted down again to resuspend in Click-iT saponin-based permeabilization and wash reagent for 15 minutes of permeabilization. After the incubation, Click-iT Plus reaction cocktail was added and incubated for 30 minutes. Cells were then washed and pelleted down for a resuspension in Click-iT saponin-based permeabilization and wash reagent containing 5 µg/ml Hoechst 33342 (Invitrogen H3570) for staining the DNA content. DNA content staining was for 15 minutes. After the DNA content staining, the cell suspensions were filtered through the cell strainer for flow cytometry. Lastly, stained cells were analyzed with flow cytometers BD LSR II and Fortessa X-20. Data were analyzed with FACSDiva and FlowJo7.

### Total RNA sequencing

From the total RNAs extracted using TRIzol (Invitrogen 15596-018), ribosomal RNAs were removed and RNA-Seq libraries were generated using TruSeq Stranded Total RNA Library Prep Gold Kit (Illumina 20020598). Total RNA-Seq libraries were sequenced as paired-end 2x100 bps using the Illumina NovaSeq platform. The sequencing reads were mapped to the human genome GRCh38 (hg38) using STAR, version 2.7.9. The mapped reads were annotated and quantified using RSEM. Differential gene expression was analyzed using R package, DESeq2. DAVID (https://david.ncifcrf.gov/) was used to perform the gene ontology analysis.

### qRT-PCR

cDNAs was synthesized from total RNAs were extracted with TRIzol, using oligo dT primers and the RevertAid First Strand cDNA Synthesis Kit (Thermo K1621) according to the manufacturer’s instructions. qPCR was then conducted using SYBR Green real-time PCR master mix (Applied Biosystems 368702). Quantification was performed by the relative standard curve method and normalized to the GAPDH. qPCR was performed using the following primers: LIN28A_Fwd 5’-TGCGGGCATCTGTAAGTGGT-3’, LIN28A_Rev 5’-GGAACCCTTCCATGTGCAGC-3’, REELIN_Fwd 5’-ATCTGCATCTGCGACGAGAG-3’, REELIN_Rev 5’-CAGTTTGCCTCGGTGACTCT-3’, GAPDH_Fwd 5’-CCACTCCTCCACCTTTGAC-3’, and GAPDH_Rev 5’-CACCCTGTTGCTGTAGCCA-3’.

### RNA half-life analysis

To analyze the mRNA decay rate in NPCs, 1.5x10^4^ cells/cm^2^ were plated, and the following day, 4μg/ml of Act. D (Sigma A9415) was added to the NPC media to block transcription. RNAs were extracted from the cells 1.5 and 3 hours after Act. D treatment. As a control, RNA representing the baseline time point was extracted from untreated NPCs. The extracted RNA was then used for qRT-PCR analysis.

### RNA immunoprecipitation

NPCs were harvested using cold DPBS and centrifuged at 3,880 g for 5 minutes at 4°C. The cell pellet was resuspended and lysed with cold buffer D (20 mM Tris-HCl pH 8.0, 0.2 mM EDTA, 100 mM KCl) supplemented with the 40 units/ul of RNase inhibitor (Takara 2311A). The cells were then sonicated using a Biorupter (amplitude 35, 1.5 minutes, 30-second pulse, 3 cycles) and centrifuged at 18,000 g for 15 minutes at 4°C. Ten percent of the total fraction volume was directly used for RNA extraction, serving as Input. The remaining supernatant was incubated with protein A/G beads (Pierce 88802), precleared with buffer D, and 5 ug of anti-LIN28A (Cell Signaling 8706), or normal rabbit IgG (Invitrogen 026102) for 1 hour at 4°C on a rotator (SeouLin Bioscience SLRM-2M). The immunoprecipitated samples were washed 4-5 times with cold buffer D and RNAs were extracted using phenol/chloroform (Sigma P1944). cDNAs were then synthesized using oligo dT and RevertAid First Strand cDNA Synthesis Kit. PCR amplification of REELIN and GAPDH was performed using the following primers: REELIN IP_Fwd 5’-ACCTACTACGTTCCGGGACA-3’, REELIN IP_Rev 5’-TCGTGGGCCATATGGTTCAC-3’. For GAPDH amplification, the same primers used in qRT-PCR were employed.

### Western blot

For total cell lysates, cell pellets were resuspended in RIPA buffer (Biosesang R2002) by pipetting and then incubated on ice for 10 minutes. The cell lysates were then centrifuged at 18,000 g 15 minutes at 4°C. For nuclear and cytoplasmic lysates, proteins were extracted using NE-PER™ Nuclear and Cytoplasmic Extraction Reagents (Thermo 78833) according to the manufacturer’s instructions. Protein concentration were determined using a Bradford assay kit (BioRad 5000006) according to the manufacturer’s instruction. A total of 10-20 μg of total cell lysates or 5-10 μg of fractionated cell lysates was separated on 10% SDS-polyacrylamide gels and transferred onto nitrocellulose (NC) membranes (Amersham 10600002). The NC membrane was then incubated in a solution of 5% skim milk (LPS solution SKI500) in PBST (0.1% Tween 20 (Bio-Basic TB0560) in PBS (VMR E404-200TABS)) for 1 hour and subsequently incubated with the primary antibody in 5% skim milk in PBST overnight at 4°C. Primary antibodies were detected using horseradish peroxidase (HRP)-conjugated secondary antibodies (Cell Signaling 7074 or 7076). The membrane was washed 5-6 times with PBS-T before developing. Protein bands were visualized with enhanced chemiluminescence (Millipore WBKLS0500) on Chemidoc XRS+ (BioRad) and iBright (Invitrogen) imaging systems. Quantitative analysis was performed using ImageJ software.

### Immunostaining

NPCs and neurons were cultured in 8-well chamber plates (SPL 30508). Neural rosettes were cultured in 4-well plates. After 1-2 washes with DPBS, cells were fixed with cold 4% formaldehyde (Pierce 28906) for 30-60 minutes at RT. Cells were then permeabilized with 0.1% Triton-X-100 (Promega H5141) in PBS for 30 minutes at RT and blocked with 3% BSA in 0.1% PBST (0.1% Tween-20 in PBS) for 1 hour at RT.

Organoids were washed 1-2 times with PBS and fixed in cold 4% formaldehyde for 30-60 minutes at RT. Then for cryoprotection, the organoids were incubated in 30% sucrose at 4°C until they settled at the bottom of the conical tube. After rinsing 2-3 times with DPBS, the organoids were transferred into block molds (Sakura 4566, Sungwon K4556) and frozen in dry ice with O.C.T. compound (Lecia 14020108926). The samples were cryosectioned into 40-um slices and placed on slide glasses (Marienfeld HSU-0810001). Then, a water-repellent barrier around samples was drawn using a hydrophobic pen (Matsunami HMA-GMP0010). After 3 washes with DPBS, the organoid slices were permeabilized and blocked using 5% donkey serum (Abcam ab7475) in 0.1% Triton X-100 in PBS for one hour at RT.

After primary antibodies were incubated overnight at 4°C, the cells were rinsed once and washed 3 times with PBST. Fluorescent dye-conjugated secondary antibodies were incubated with the cells for 1 hour at RT. Following 5-6 washes with PBST, cells were incubated with DAPI (Sigma D9542) for 10 minutes at RT and mounted on slides using fluorescence mounting medium (DAKO S3023). Images were acquired using a confocal microscopy (Zeiss LSM800 or LSM980).

### Lentivirus (LV) production

The LV plasmid of interest and LV packaging plasmids (MDL, RSV-Rev, and VSVG) were delivered using polyethylenimine (PEI; Polysciences 23966) into HEK293T cells, grown in DMEM (Welgene LM001-07) supplemented with 10% fetal bovine serum (Young-in Frontier US-FBS-500, Gibco 12483-020). Cells were plated a day before transfection, and the media was changed 4 hours post-transfection. Culture media for LV production were harvested on day 3. LVs were concentrated by ultracentrifugation.

### Neuronal morphology analysis

To identify the morphology of a single neuron, lentivirus-expressing GFP was transduced with low MOI (MOI = ∼0.1). After taking images, the neurites of neurons expressing GFP were traced manually and neuronal morphology was analyzed. Sholl analysis and neurite tracing was performed using Simple Neurite Tracer (SNT) software from Fiji.

### Multielectrode array (MEA)

The 96-well MEA plate from Axion Biosystems was prepared by coating with 500 ng/ul of PLO and 5 μg/μl of Lam. Then, 10,000 NPCs were plated in each well and differentiated into neurons as previously described. Culture media was changed every 2-3 days, with measurements taken before each media changes. Recordings were performed using the Maestro MEA system and AxIS software (Axion Biosystems). The MEA plate was allowed to stabilize for 15 minutes in the Maestro Instrument before data recording for an additional 15 minutes to calculate the spike rate per well. Multielectrode data analysis was performed using the default settings in the Axion Biosystems Neural Metrics Tool.

### ChIP sequencing and chromatin enrichment score analysis

One million neural cells were collected and washed 1-2 times with PBS and then fixed in cold 1% formaldehyde in PBS for 10 minutes at RT. Fixation was stopped by quenching with 0.125 M glycine for 5 minutes at RT. Protease inhibitor cocktails were added to the harvested cells, which were then sonicated in a microtube AFA fiber snap-cap (Covaris 520045) using a Covaris S220 (peak power 140, duty factor 10, cycles per burst 200, time 55 seconds) until DNA fragments ranged from 200 to 700 bp. Sheared chromatins were incubated with AGO1 primary antibodies (WAKO 015-22411) overnight at 4°C on the rotator. To precipitate chromatin-antibody complexes, samples were incubated with protein A/G Dynabeads for 2 hours at 4°C on a rotator (Nutator mixer KA.PM-6249following an overnight non-specific blocking of the Dynabeads with 0.1% BSA at 4°C. Chromatin-bound Dynabeads were washed twice with cold RIPA-LS (0.1% SDS, 1% Triton X-100, 1 mM EDTA, 10 mM Tris-HCl pH 8, 140 mM NaCl, 0.1% Sodium deoxycholate), twice with RIPA-HS (0.1% SDS, 1% Triton X-100, 1 mM EDTA, 10 mM Tris-HCl pH 8, 500mM NaCl, 0.1% Sodium deoxycholate), twice with cold RIPA-LiCl (0.25 M LiCl, 0.5% NP40, 1 mM EDTA, 10 mM Tris-HCl pH 8, 0.5% Sodium deoxycholate), and once with cold 10 mM Tris pH 8 buffer. Samples were incubated for 10 min at 37°C to tagmentation with lab-made Tn5 transposase. Samples were washed twice with cold RIPA-LS and twice with cold TE buffer (10 mM Tris-HCl, pH 8, 1 mM EDTA). They were then incubated for 1 hour at 55°C, followed by 10 hours at 65°C for de-crosslinking. The immunoprecipitated DNA was eluted using the MinElute PCR Purification Kit (QIAGEN, 28004). Input DNA prepared with 1% sonicated lysate was performed decrosslinking followed by tagmentation. For ChIP sequencing, the libraries were sequenced using an Illumina NovaSeq6000 SE50. The ChIP sequencing data were mapped to the human genome GRCh38 (hg38) using Bowtie 2.4.1. AGO1 peaks in NPCs were called using MACS2 with option -p 0.005. The mapped bam files were converted into bedGraph files using MACS2 and normalized using total read counts. The bedGraph files were converted into bigWig files using bedGraphToBigWig.

To calculate the enrichment score, peak regions were initially defined using the aggregate option in bwtool package. The positions −1 bp and +1 bp flanking the boundaries of each peak were defined for normalization. The average peak intensity was calculated by dividing the total peak intensity across the annotated peak regions by the length of the peak region. This average peak intensity was then normalized by the average intensity at the −1 bp and +1 bp boundary positions to derive the enrichment score. A minimum average peak intensity threshold of 500 was established for analysis.

### Assessment of AGO1 and LIN28A expression using single cell atlas of the human brain

To elucidate the cell type and developmental stage-specific expression patterns of AGO1 and LIN28A, comprehensive single cell atlas of the developing human brain was utilized^77^. The atlas spans from 7 gestational weeks to 90 years of age, with 41 clusters annotated as 10 major cell types and 22 cell subtypes. In order to comprehensively understand the representation of stages across a wider range of contexts, the stages presented in the source data were categorized into 5 larger categories: fetal, neonatal, childhood, adolescence, and adult. Similarly, the cell types were classified into 5 major groups: glial (astrocyte, microglia, oligodendrocyte, OPC), NPC (RG, neuroblast), neuronal (excitatory neurons, inhibitory neurons), vascular (endothelial), and others.

### Trajectory analysis using NPC subset data from single cell atlas

The temporal expression dynamics of AGO1 and LIN28A within the fetal NPC cells (C3, C4, C11, C18, C24, C26, C33, C37) of the single-cell atlas were analyzed. The subset dataset was reprocessed for the trajectory analysis. Normalization and log transformation of raw counts were conducted with Scanpy^78^ (v.1.8.2). Scvi-tools^79^ (v.1.0.3) was used to adjust for batch effect between samples, with ‘SampleID’ as batch key and ‘Dataset,’ ‘Assay,’ and ‘Library’ as categorical covariate key. Five thousand highly variable genes were selected for this process.

A pseudotime trajectory analysis was conducted using Palantir^80^. Diffusion map was computed with scvi-corrected embeddings. One of the cells with the youngest age was chosen as the starting cell for the trajectory, whereas the terminal cells were automatically identified by Palantir. Default parameters were used for all processes. Gene expression trends for each trajectory were computed with the compute_gene_trends function. Expression patterns of AGO1 and LIN28A within each trajectory lineage were visualized using the plot_gene_trends function.

### LIN28A overexpression and knockdown in NPCs

For LIN28A overexpression, NPCs were transfected with LIN28A-expressing plasmids using lipofectamine (Thermo STEM00003). The pcDNA3(+)-FLAG-LIN28A plasmid, a gift from Dr. V. Narry Kim (Seoul National University, Republic of Korea), was used. The pcDNA3(+) null vector served as the control. After 24 hours, cells were collected for further experiments.

For LIN28A suppression, LV-expressing shRNAs were utilized. Two different shRNAs targeting LIN28A (shLIN28A#1 (TRCN0000021802) and shLIN28#2 (TRCN0000021803) were designed by the RNAi Consortium of the Broad Institute and cloned into the lentiviral vector pLKO.1.5-puro. Non-targeted shRNAs with scrambled sequences, shSCR#1 and shSCR #2, were used as controls. The pLKO.1-Scrambled and pLKO.1 -TRC control vectors were gifts from Anthony Leug (Addgene 136035)^81^and David Root (Addgene 10879)^82^, respectively. NPCs were infected with the LVs expressing shRNAs (MOI=∼0.5). Following a 24-hour incubation, the cells underwent antibiotic selection in media containing 1 µg/ml puromycin. Samples were collected 72 hours after LV transduction for further experiments.

### Analyzing neural rosette circularity

After differentiating hESCs into neural rosettes, the rosettes were stained with an anti-ZO-1 antibody (Thermo 61-7300). Images were then acquired using confocal microscopy. The longest and shortest axes of the lumens within the neural rosettes were arbitrarily determined and measured. Circularity was defined as the ratio between the lengths of the longest and shortest axes.

To analyze the effect of REELIN on neural rosette circularity, 200 ng/mL of recombinant human REELIN (R&D Systems 8546-MR-50) was added to the culture media starting from the day the ESCs were transferred to non-adherent dishes. Fresh REELIN was supplemented every 2 days, concurrent with media changes. The control group was treated with PBS. REELIN treatment was applied for 7-10 days during the EB floating culture phase and continued for 3-4 days following plating on Matrigel. REELIN treatment was maintained until fixation to assess rosette circularity.

To investigate the effect of LIN28A on neural rosette circularity, LIN28A expression was suppressed in *AGO1* KO ESCs using shRNAs. *AGO1* KO hESCs were dissociated into single cells and plated on a Matrigel-coated plate using mTeSR with FGF2 and Y27632. One day after plating, the hESCs were transduced with LVs expressing shRNAs (shSCR#1, #2 and shLIN28A#1, #2) along with puromycin-resistant genes (MOI=∼0.5). After 24 hours of LV infection, the cells were treated with 2 μg/ml puromycin for 48 hours. The surviving hESCs were allowed to grow and form colonies. LIN28A knockdown in these colonies was confirmed through qRT-PCR and Western blot analyses. Selected *AGO1* KO hESC clones with LV infection were then used to analyze neural rosette circularity.

### Data presentation and statistical analysis

Representative experiments were repeated in at least 3 independent biological replicates or organoids, yielding similar results, Data are presented as mean ± SEM. NGS datasets underwent normalization and significance correction using R and python tools such as DESeq2, Bowtie2, MACS2, Scanpy, Scvi-tool and Palantir with adjusted p values (padj) corrected for multiple testing using false discovery rates (FDR). Statistical analyses were conducted using unpaired Student’s t-tests, r multiple t-test, one-way ANOVA with multiple comparisons with Tukey’s or Sidak’s test, or two-way ANOVA with repeated measures or Sidak’s test in Prism (GraphPad). Significance is indicated in the figures as non-significant (ns) p>0.05, *p < 0.05; **p < 0.01; ***p < 0.001; and ****p < 0.0001.

## Acknowledgments and funding sources

We thank members of our laboratory for critical reading of the manuscript, Mary Lynn Gage for editorial comments, and Yeseul Kang for invaluable help in creating the description figures and graphic abstracts. This work was supported by the National Research Foundation of Korea (NRF) (2022R1A2C3003115 to D. L.; RS-2024-00439474 to J.-Y. A.; RS-2024-00335144, RS-2024-00398786, RS-2024-00407383 to J.H.), and the Institute for Basic Science (IBS-R002-A2 to S.-H.L., IBS-R002-A1 to J.H.) from the Ministry of Science and ICT of Korea. F.H.G was supported by the National Institutes of Health of USA (R01 AG056306, UCSD - R01 AG057706, and UCSD - R01 AG056511), the American Heart Association and the Paul G. Allen Frontiers Group (Grant #19PABHI34610000/TEAM LEADER: Fred H. Gage/2019), The JPB Foundation and the Milky Way Research Foundation. We also acknowledge the support by the NGS Core Facility of the Salk Institute with funding from NIH-NCI CCSG: P30 014195. This research was also supported by Ph.D. Fellowship Program through NRF funded by the Ministry of Education (2018H1A2A1059772 to H. D) and Basic Science Research Program (2022R1A6A3A13073152 to D. K, RS-2024-00410041 to I. A).

