## supplementary figures for "AGO1 in neural progenitor cells orchestrates brain development and sociability via LIN28A-REELIN axis"

### Do et al. Extended Data Fig. 1

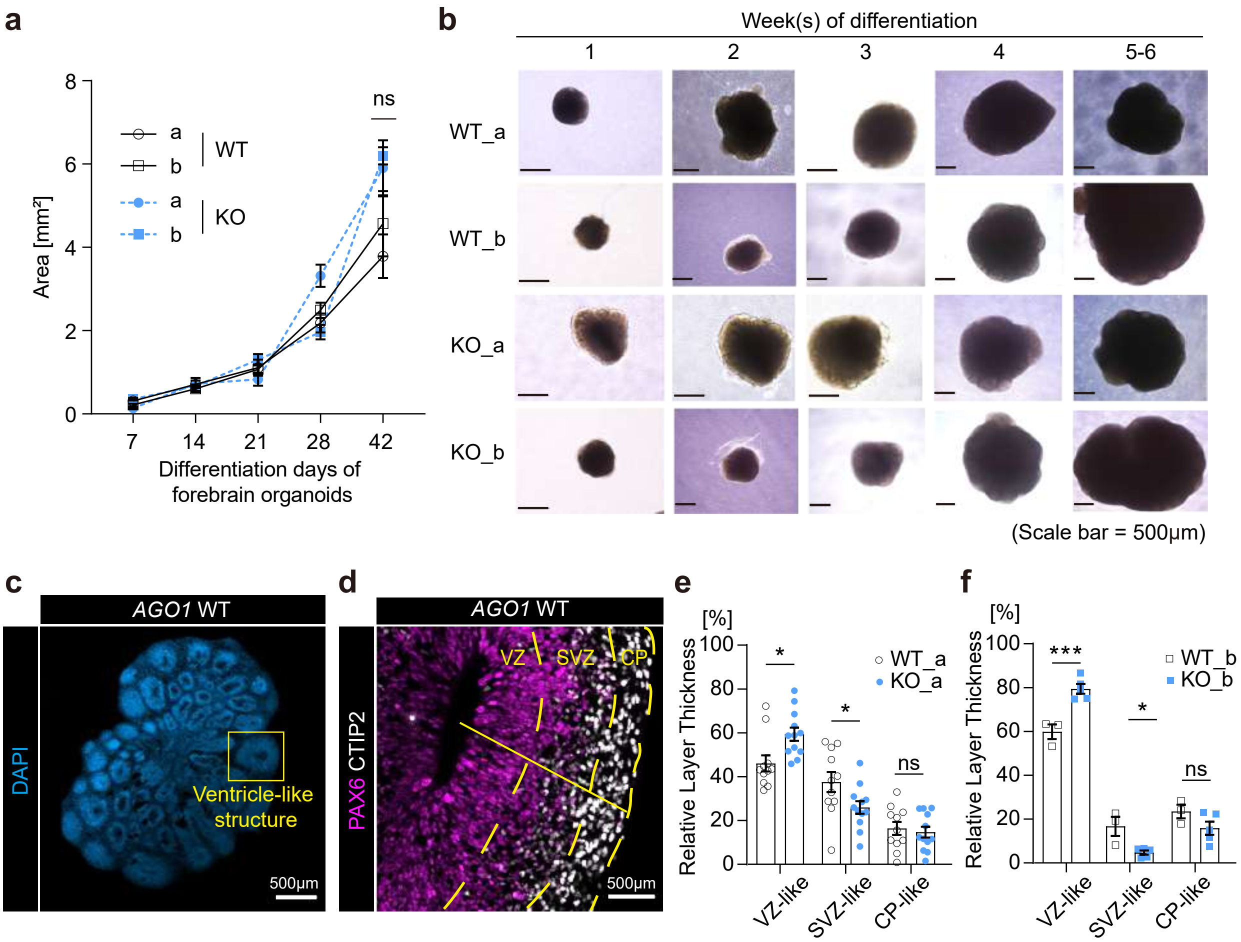

### Do et al. Extended Data Fig. 2

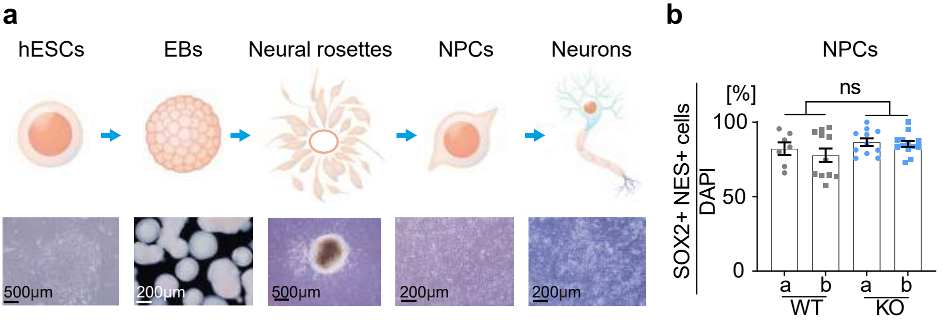

### Do et al. Extended Data Fig.3

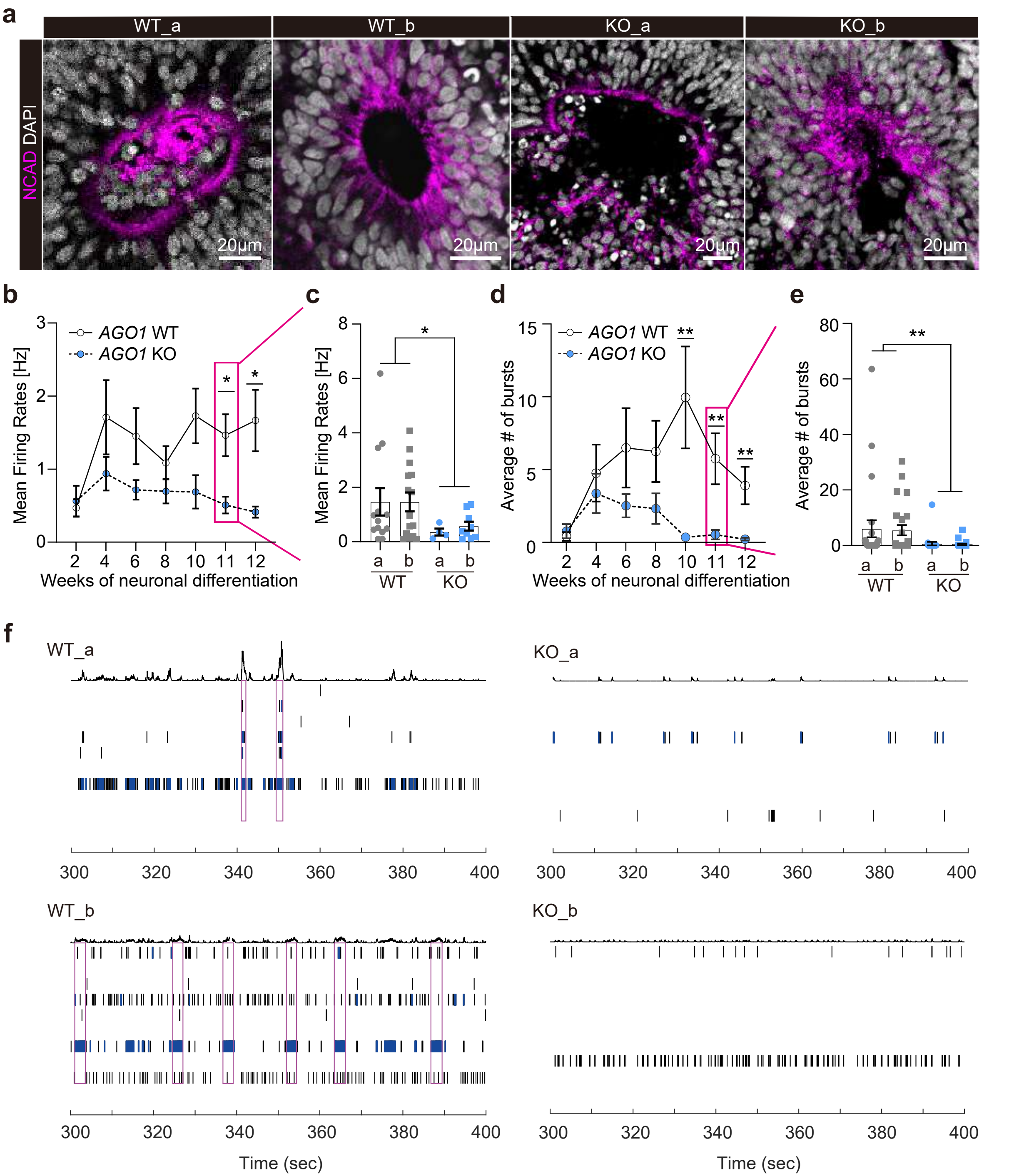

### Do et al. Extended Data Fig. 4

**a**

AGO1 ChIP-seq in NPCs

| Annotated region | Numbers of peaks |
|---|---|
| Intergenic | 11785 |
| Intron | 10861 |
| Promoter-TSS | 1290 |
| TTS | 330 |
| 3'UTR | 219 |
| Non-coding | 130 |
| Exon | 119 |
| 5'UTR | 47 |
| Total | 24781 |

**b**

Top 5 up- and down-regulated DEGs with fold change in AGO1 KO NPCs

| | Gene symbol | Fold change | padj |
|---|---|---|---|
| UP regulated genes | TAL2 | 2.77 | 2.02E-02 |
| | ADAMTS20 | 2.76 | 7.35E-05 |
| | LIN28A | 2.29 | 5.55E-16 |
| | S100A6 | 2.17 | 1.77E-04 |
| | NRN1 | 2.15 | 5.25E-15 |
| DOWN regulated genes | RELN | -4.39 | 1.49E-08 |
| | NKX6-1 | -4.00 | 1.43E-05 |
| | SHOX2 | -3.96 | 2.10E-28 |
| | HOXB4 | -3.80 | 3.45E-03 |
| | HOXB3 | -3.56 | 9.49E-07 |

**c**

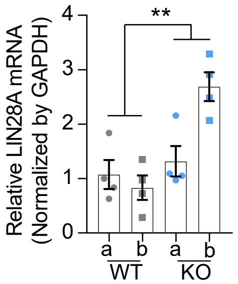


**d**

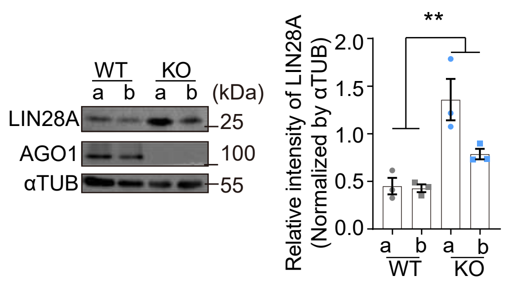


**e**

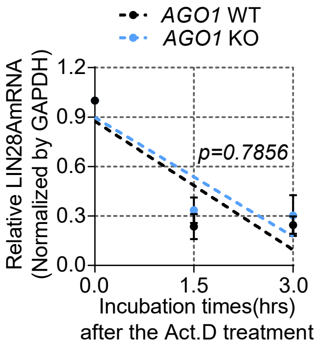


**f**

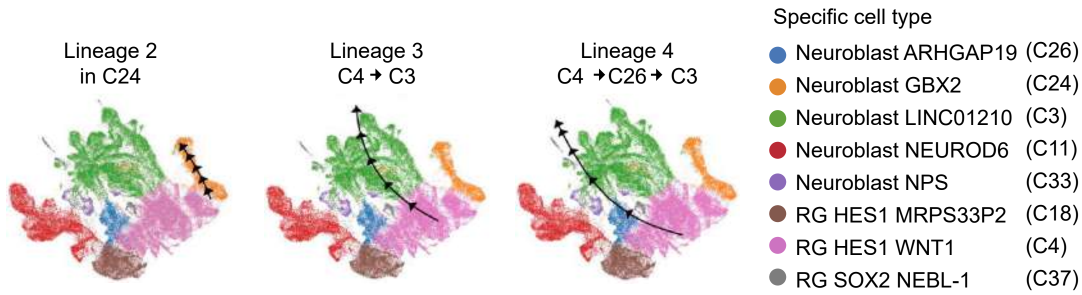


**g**

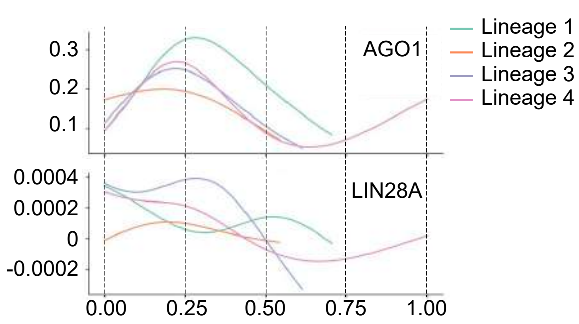


**h**

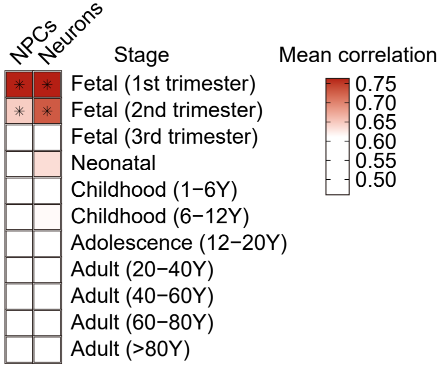

Do et al. Extended Data Fig. 5

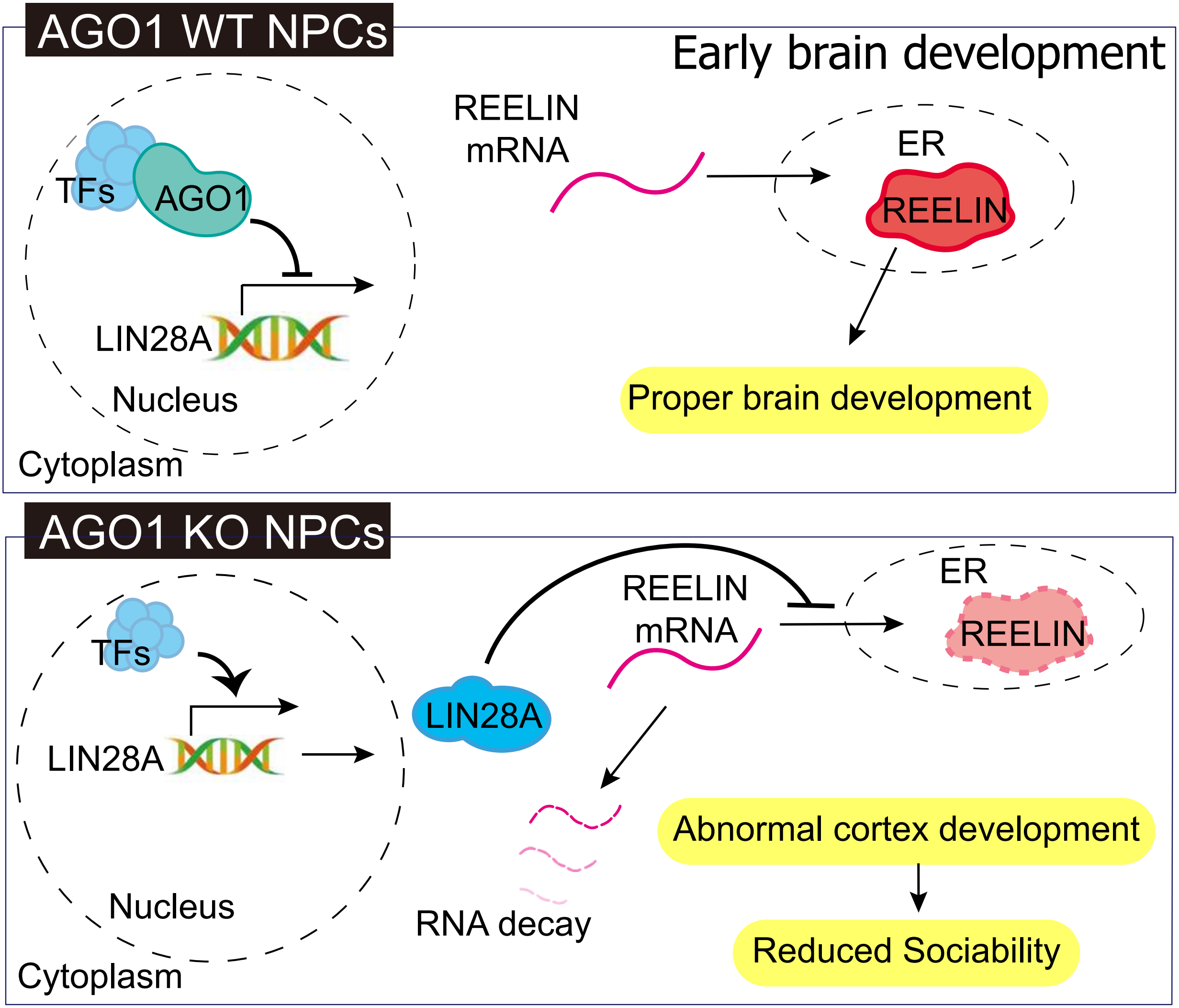
